# Inferring immunological control mechanisms from TKI dose alterations in CML patients

**DOI:** 10.1101/722546

**Authors:** Tom Hähnel, Christoph Baldow, Artur C. Fassoni, Joëlle Guilhot, François Guilhot, Susanne Saussele, Satu Mustjoki, Stefanie Jilg, Philipp J. Jost, Stephanie Dulucq, François-Xavier Mahon, Ingo Roeder, Ingmar Glauche

## Abstract

Recent clinical findings in chronic myeloid leukemia (CML) patients suggest that the risk of molecular recurrence after stopping tyrosine kinase inhibitors (TKI) treatment substantially depend on an individual, leukemia-specific immune response. However, it is still not possible to prospectively identify patients that will most likely remain in a long-term treatment free remission (TFR). Here, we use a mathematical model for CML, which explicitly includes an anti-leukemic (presumably immunological) effect and apply it to a set of patients (n=60) for whom BCR-ABL/ABL time courses had been quantified before and after TKI stop. We demonstrate that such a feedback control is conceptually necessary to explain long-term remission as observed in about half of the patients. Based on simulation results we classify the patient data sets into three different groups according to their predicted immune system configuration. While one class of patients requires a complete CML eradication to achieve TFR, other patients are able to control the leukemia after treatment cessation. Among them, we identified a third class of patients, which only maintains TFR if an optimal balance between leukemia abundance and immunological activation is achieved before treatment cessation. Further, we demonstrate that the immune response classification of the patients cannot be obtained solely from BCR-ABL measurements *before* treatment cessation. However, our results strongly suggest that changes in the BCR-ABL dynamics arising after system perturbations, such as TKI dose reduction, holds the information to predict the individual outcome after treatment cessation.

## Introduction

Chronic myeloid leukemia (CML) is a myeloproliferative disorder, which is characterized by the unregulated proliferation of immature myeloid cells in the bone marrow. CML is caused by a chromosomal translocation between chromosomes 9 and 22. The resulting *BCR-ABL* fusion protein acts as constitutively activated tyrosine kinase triggering a cascade of protein phosphorylation, which deregulate cell cycle, apoptosis regulation, cell adhesion and genetic stability. Due to their unregulated growth and their distorted differentiation, immature leukemic cells accumulate and impair normal hematopoiesis in the bone marrow, leading to the patient’s death if left untreated.

Tyrosine kinase inhibitors (TKIs) specifically target the kinase activity of the BCR-ABL protein with high efficiency and have been established as the first line treatment for CML patients (1). Individual treatment responses are monitored by measuring the proportion of BCR-ABL transcripts relative to a reference gene, e.g. ABL or GUS, in blood cell samples by using quantitative real-time polymerase chain reaction (qRT-PCR) (2–4). Most patients show a typical bi-exponential treatment response with a rapid, initial decline (α slope), followed by a moderate, second decline (β slope) (5–7). Whereas the initial decline can be attributed to the eradication of proliferating leukemic cells, the second decline has been suggested to result from a slower eradication of quiescent leukemic stem cells (3,4,8,9). Within five years of treatment, about two thirds of the patients achieve a major molecular remission (MMR), i.e. a BCR-ABL reduction of three logs from the baseline (MR3), while at least one third of these additionally achieve a deep molecular remission (DMR, i.e. MR4 or lower) (4,7,10).

TKI discontinuation has been established as an experimental option for well responding patients with DMR for at least one year. Different studies independently confirmed that about half of the patients show a molecular recurrence, while the others stay in sustained treatment-free remission (TFR) after TKI stop. Consistently, most patients present with a recurrence within 6 months, while only a few cases are observed thereafter (11–15). The overall good response of those patients after restarting treatment with the previously administered TKI indicates that clonal transformation and resistance occurrence is not a primary problem in CML. As it appears unlikely that even a sustained remission truly indicates a complete eradication of the leukemic cells, other factors have to account for a continuing control of a minimal, potentially undetectable residual leukemic load. Although treatment discontinuation is highly desirable to reduce treatment-related side-effects and lower financial expenditures (16–19), it is still not possible to prospectively identify those patients that are at risk for a molecular recurrence. Investigations of clinical markers and scores to predict the recurrence behavior of patients after the treatment cessation revealed that both TKI treatment duration and the duration of a DMR were also associated with a higher probability of TFR (11,15,20,21). However, it is still unclear whether the dynamics of the initial TKI treatment response (e.g. the initial slope of decline) correlate with the remission occurrence after treatment discontinuation.

The underlying mechanisms of the recurrence behavior after TKI stop are still controversial. While fewer recurrences for patients with longer treatment suggest that a leukemic stem cell exhaustion is an important determinant, it is not a sufficient criteria to prospectively identify non-recurring patients (11,20). Favorable outcomes of treatment discontinuation for patients that were previously treated with immune-modulating drugs, such as IFN*-*α, suggest that immunological factors might play an additional and important role (15,21,22). In this context, it has been demonstrated that specific subpopulations of dendritic cells and natural killer cells, as well as the cytokine secretion rate of natural killer cells are associated with higher probabilities of a treatment-free remission (23,24). Furthermore, there are several reports about patients with low but detectable BCR-ABL levels over longer time periods after therapy discontinuation that do not relapse (12,25). This is a strong indicator that also other control mechanisms, such as the patient’s immune response, are important determinants of a TFR.

Mathematical models have propelled the conceptual understanding of CML treatment dynamics (5–7,9,26–30). Especially the long-term effect of TKI treatment on residual stem cell numbers and the effect of combination therapies were in focus. In a recent publication, we provided evidence that TKI dose reduction is a safe strategy for many patients in sustained remission while preserving the anti-leukemic effect (9). Complementary efforts also accounted for interactions between leukemic and immune cells (31–35). In a prominent approach, Clapp et al. used a CML-immune interaction to explain fluctuations of BCR-ABL transcripts in TKI-treated CML patients (33). However, it remains elusive to which extend an immunological control is a crucial mediator to distinguish patients that maintain TFR from those that will eventually relapse.

Here, we used BCR-ABL time courses of TKI-treated CML patients that were enrolled in previously published TKI discontinuation studies from different centers in Europe. In particular, we focused on patients for which *complete time courses* during the initial TKI therapy *and* after treatment cessation are available. Therefore, potential correlations between response dynamics, remission occurrences and timings after cessation become accessible. We demonstrate that the residual disease level at the time of the TKI stop are not sufficient to predict the molecular recurrence after treatment discontinuation. Instead, we use a mathematical model of TKI-treated CML, which includes a patient-specific, CML dependent immune component and demonstrate that three different immunological configurations can determine the overall outcome after treatment cessation. We further investigate how this patient-specific configuration can be estimated from system perturbations, such as TKI dose reduction scenarios prior to treatment cessation.

## Methods

### Patient selection

We used time courses of 60 TKI-treated CML patients, for whom TKI-therapy had been stopped as a clinical intervention. Informed written consent was obtained from each subject according to the local regulations of the participating centers. Detailed information on the patient cohort is available in the Supplement Materials. For all these patients, serial BCR-ABL/ABL measurements before as well as after cessation are available. For the purpose of this analysis, the date of a molecular recurrence after cessation was defined as the first detected BCR-ABL/ABL -ratio above 0.1%, indicating a loss of MR3, or the re-initiation of TKI treatment, whatever was reported first.

Furthermore, we selected patients, which received TKI monotherapy before stopping, which have been monitored at a sufficient number of time points to estimate the initial and secondary slopes and which present with the typical bi-exponential response dynamic (Fig. 1). The 21 selected patients, fulfilling those criteria, were compared with the full patient cohort (n=60) and showed no obvious differences for the initial BCR-ABL levels, treatment duration, recurrence behavior, follow up duration, recurrence times and used TKI, and are, therefore, considered to be representative examples. (Fig. S1). Moreover, the overall recurrence behavior of the selected patient cohort is comparable to larger clinical studies (12,15).

**Fig. 1:**
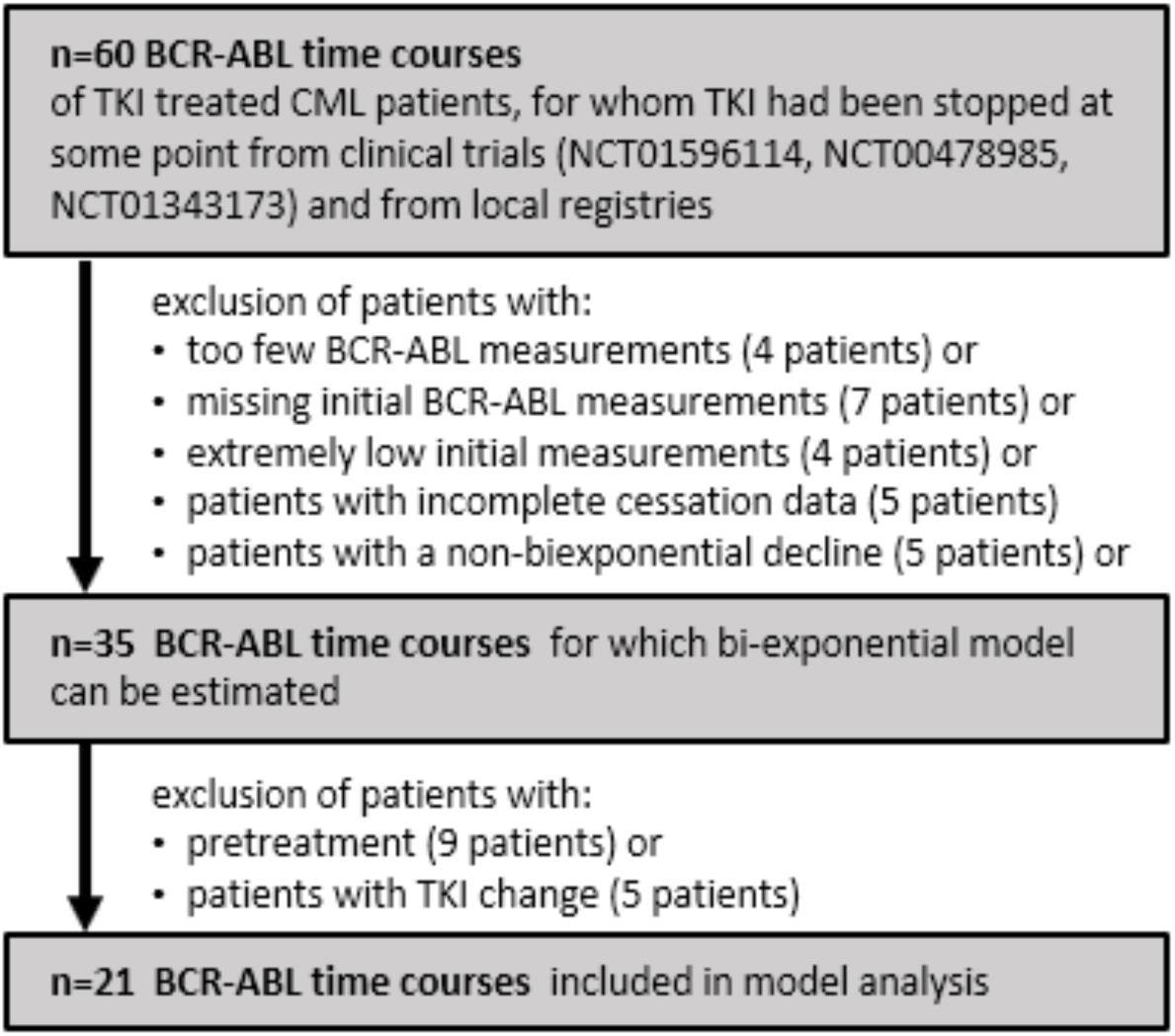
Strategy for patient selection. We excluded patients with less than 5 BCR-ABL measurements during TKI treatment, missing or extremely low initial measurements (i.e., first measurement is only available after more than 10 months or below MMR). Furthermore, we only included patients with a biexponential decline in which the initial slope was steeper than the second slope. We also only selected patients that were under continuous therapy with one TKI, thereby excluding patients with a pre-treatment, TKI-change during the treatment, combination therapy and missing therapy/cessation information.

### Bi-exponential parameterization of treatment response

For a formal analysis, we describe the time dependent BCR-ABL/ABL-ratio of TKI-treated patients on the log-scale using the logarithm of a bi-exponential function, i.e. log_10_((*BCR* – *ABL*) / *ABL=LRATIO*(*t*)=log_10_(*Ae*^*αt*^*+Be*^*βt*^). Each patient’s BCR-ABL dynamics is, therefore, defined by the parameters A, B, α and β, which are estimated by a maximum likelihood (ML) approach (*R* version 3.4.4, package *maxLik* version 1.3.4) as described in (36). To obtain uniquely identifiable solutions in the ML procedure, we restrict *α*< *β*, according to the selection of patients showing a rapid initial decline *α* compared to a slower long-term decay *β*.

### Mathematical model of TKI-treated CML

For our analysis, we apply an ordinary differential equation (ODE) model, which we proposed earlier in a methodological article qualitatively comparing a set of CML models with different functional interaction terms between leukemic cells and immune cells (35).

This model is formally described by:

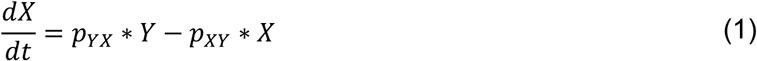

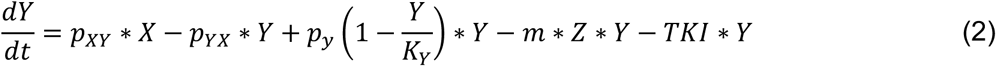

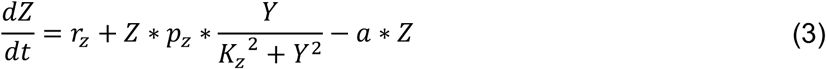

In brief, the ODE model distinguishes between a population of quiescent leukemic cells (*X*) and a population of actively cycling leukemic cells (*Y*), which proliferate with the rate *p*_*Y*_, whereas the growth is limited by a carrying capacity *K*_*y*_. Leukemic cells can switch reversibly between the active and the quiescent state with transition rates *p*_*XY*_ and *p*_*XY*_. TKI treatment is modelled by a kill rate *TKI* =*TKI*_0_, which acts on proliferating cells *Y*, but does not affect quiescent cells *X*. A complete eradication of leukemic cells is defined as a decrease of leukemic cells in *X* and *Y* below the threshold of one cell. The corresponding BCR-ABL/ABL ratio in the peripheral blood are calculated as the ratio of proliferating leukemic cells to the carrying capacity *K*_*Y*_ (see Supplementary Material for details).

Furthermore, the model integrates a population of CML-specific immune effector cells (*Z*), which are generated at a constant, low production rate *r*_*z*_ and undergo apoptosis with rate *a*. They eliminate proliferating leukemic cells *Y* with the kill rate *m*. The leukemia-dependent recruitment of immune cells follows a nonlinear functional response where *p*_*z*_ and *K*_*z*_ are positive constants. This functional response leads to an *optimal* immune cell recruitment for intermediate leukemic cell levels, i.e.: for low numbers of proliferating leukemic cells (*Y* < *K*_*z*_), the immune cell recruitment increases and the immune cells *Y* are stimulated to replicate in presence of proliferating leukemic cells *Y*, reaching a maximum *p*_*z*_/(2*K*_*Z*_) when *Y* = *K*_*Z*_. For higher leukemic cell numbers (*Y* > *K*_*Z*_) the immune cell recruitment decreases with *Y*, reflecting the assumption that the proliferation of immune cells is decreased for high levels of proliferating leukemic cells *Y*. This assumption follows recent findings, suggesting that a high load of CML cells inhibits the immune effector cells’ function and number (37). As a result we obtain an *immune window* (defined by the interval 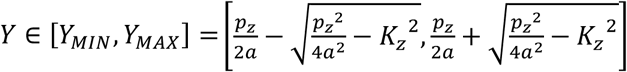) for which the recruitment exceeds the degradation rate *a* of the immune cells and leads to an optimal immune response (see Supplementary text).

For all patients, we use fixed values for the immune mediated killing rate *m*, the proliferation rate *p*_*Y*_, the carrying capacity *K*_*Y*_, the immune cells natural influx *r*_*z*_ and the immune cells apoptosis rate *a*. In contrast, the transition rates *p*_*XY*_ and *p*_*YX*_, the TKI kill rate *TKI* and the immune parameters *k*_*z*_ and *p*_*z*_ are estimated with different strategies using an *Approximate Bayesian Computation* approach (see Supplementary Material).

As a control (i.e. null model), we use a reduced version of the outlined ODE model *without* an immunological component. Herein, all the corresponding parameters values are set to zero (*p*_*z*_ = *K*_*z*_ = *a* = *r*_*z*_ = *m* = 0).

## Results

### Initial TKI response dynamics show no difference between recurring and non-recurring patients

We investigate whether the dynamics of TKI treatment response correlated with disease recurrence. Therefore, we analyzed the BCR-ABL dynamics of individual patients and fitted a bi-exponential model to each time course. We obtained estimates of the slopes α and β, describing the initial and long-term decline, respectively (Fig. 2A/B, Fig. S2). We compared the slopes between recurring and non-recurring patients (Fig. 2C/D), assessed treatment duration (Fig. 2E), time in MR4.5 (4.5 log reduction from baseline, Fig. 2F) and the estimated BCR-ABL levels at cessation time (Fig. 2G). We observed no significant differences between recurring and non-recurring patients for any these values. Furthermore, we did not find a particular co-occurrence between the parameters α and β, which would have allowed to identify distinct clusters for recurring and non-recurring patients (Fig. 2H). Although our conclusions are based on a limited amount of data, we did not detect obvious differences between recurring and non-recurring patient groups, which could potentially serve as a predictive measure to prospectively identify CML patients that show a molecular recurrence after treatment cessation.

**Fig. 2.**
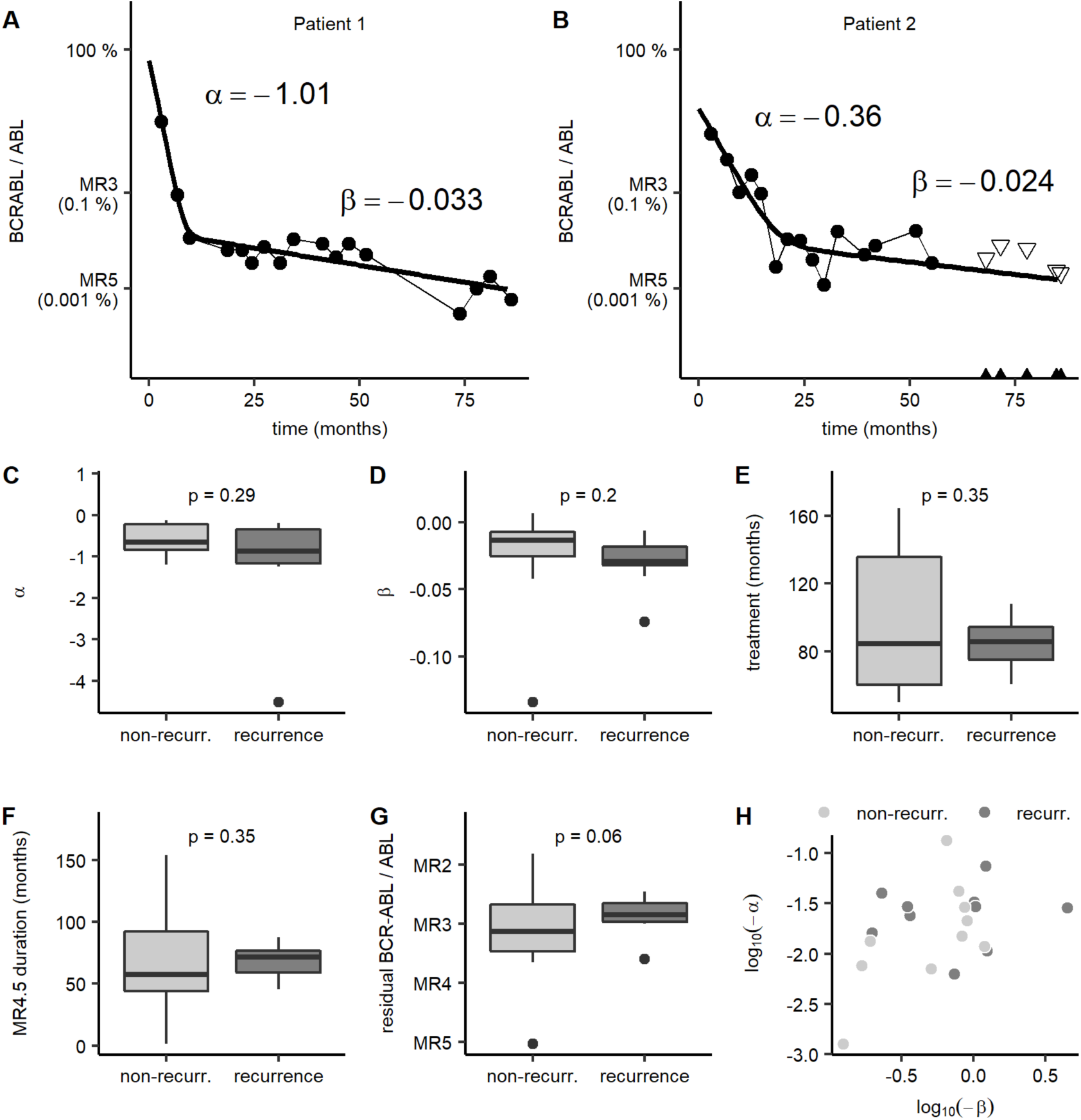
No differences of clinical data between recurring and non-recurring patients. **A/B**: Clinical data and bi-exponential fit with corresponding α and β values for two representative patients. BCR-ABL measurements are shown as dots. Triangles indicate the quantification limit (QL) for undetectable BCR-ABL levels, derived from the measurement of the reference gene ABL (*QL* = 3/*ABL*) according to the laboratory recommendations. The corresponding bi-exponential fit (thick line) consists of the initial fast decline α and the second, slow decline β. Only data until TKI cessation is shown. **C-H**: Comparison of recurring (dark grey) and non-recurring (light grey) patients by analyzing the clinical data and BCR-ABL time courses before treatment stop. P-values of the corresponding Kolmogorov-Smirnov tests are shown. **C**: Steepness of the first, fast decline (α). **D**: Steepness of the second, slow decline (β). **E**: Treatment duration before cessation. **F**: Time of deep molecular remission (DMR, reduction of 4.5 logs from the baseline) while treated, **G**: BCR-ABL levels at cessation estimated using the corresponding individual bi-exponential fits. **H**: Distribution of α and beta for recurring and non-recurring patients.

### Individual BCR-ABL dynamics after TKI stop can be explained by a patient-specific immune component

Next, we raise the question whether a simple mathematical model of leukemic treatment response is sufficient to describe the BCR-ABL data before *and* after treatment stop. As a control (i.e. a *null* model), we use a reduced version of the suggested ODE model *without* an immunological component. In this model, leukemic stem cells can reversibly switch between the quiescent (*X*) and proliferating (*Y*) state with corresponding transition rates *p*_*XY*_ and *p*_*YX*_. The TKI treatment has a cytotoxic effect on proliferating cells in *Y* (Fig. 3A, see Methods). This model is adapted to each individual patient time course by estimating the specific model parameters from the pre-cessation BCR-ABL data. Due to the bi-exponential BCR-ABL decline, the reduced model *without* an immunological component predicts a complete eradication of residual disease levels only for very long treatment times. Thus, treatment cessation at any earlier time point will eventually lead to recurrence (Fig. 3B/C, Fig. 3G/H), which clearly opposes clinical findings (11–15).

**Fig. 3:**
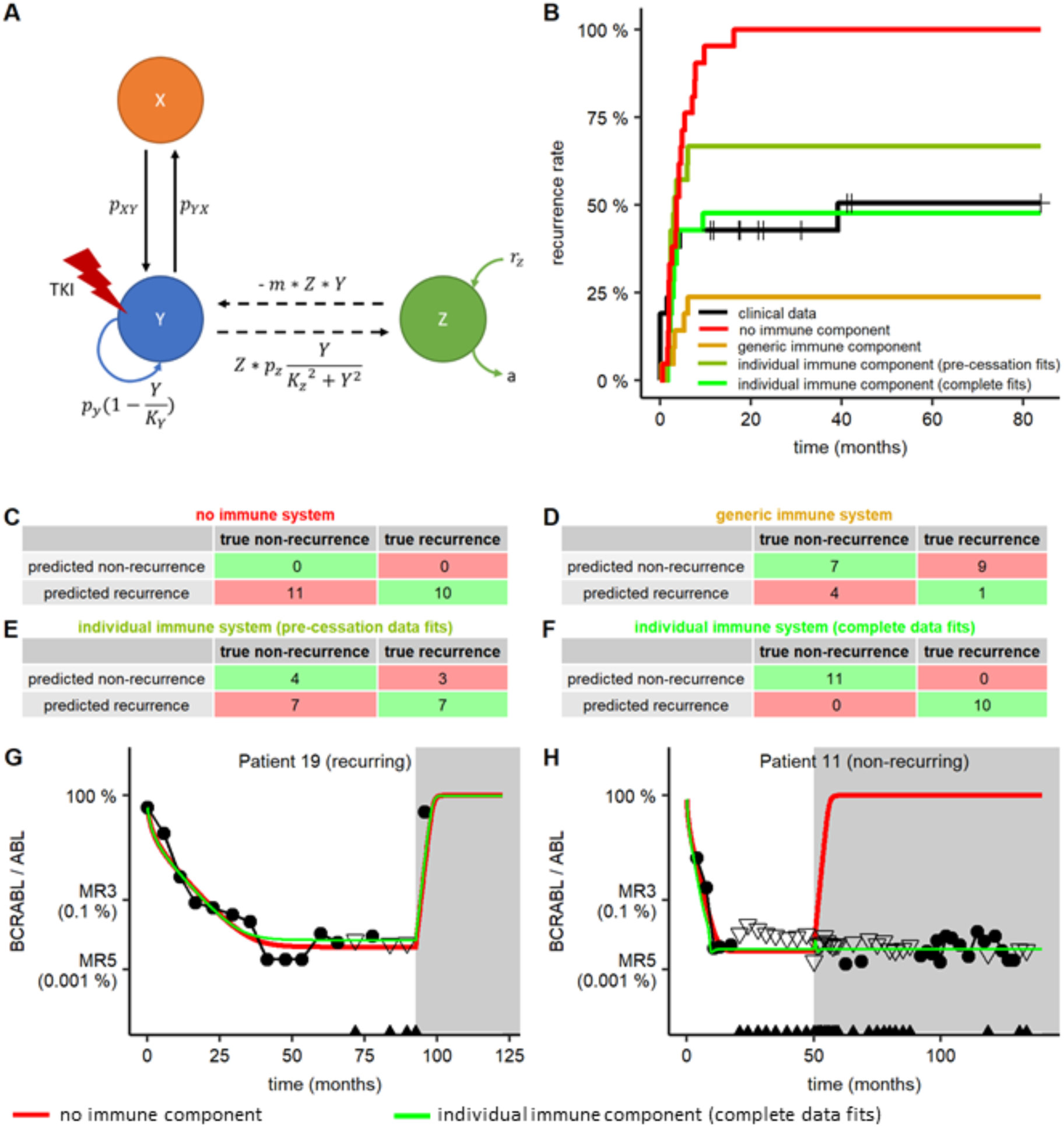
Model comparison. **A**: General scheme of the model setup indicating the relevant cell populations and the rates that govern their dynamical responses. **B**: Kaplan-Meier estimators comparing the cumulative recurrence rates of four different models with the clinical data (black): the reduced model *without* an immune component (red, C), and three different configuration of immune model (D: generic immune system configuration (fitting strategy I – brown); E: individual immune system configuration estimated by fitting pre-cessation data of the BCR-ABL time courses (fitting strategy II – olive-green); F: individual immune system configuration estimated by fitting the complete BCR-ABL time courses (fitting strategy III – green)). **C-F**: Corresponding contingency tables comparing individual recurrence predictions for the four model configurations (table headings are colored accordingly for subfigure B). **G/H**: Examples of clinical data for a representative recurring (G) and a non-recurring (H) patient with corresponding model predictions for the reduced model *without* an immune component (red) and the immune system model using an individual immune system configuration estimated by fitting complete BCR-ABL time (fitting strategy III – green). BCR-ABL measurements are shown as black dots. Black triangles indicate the lower quantification limit for undetectable BCR-ABL levels. The grey area indicates the time after treatment cessation.

Clinical studies suggest that immunological components can potentially control minimal residual disease levels and, therefore, might prevent (molecular) recurrences after TKI stop (23,24). Therefore, we use an extended ODE model that explicitly considers an immune component (35). In brief, immune cells in *Z* have a cytotoxic effect on proliferating leukemic cells in *Y*. The proliferation of immune cells is stimulated in the presence of peripheral, proliferating leukemic cells in *Y* by an immune proliferation rate *p*_*z*_. High leukemic cell levels suppress an efficient immune response by the immune inhibition constant *K*_*z*_ (Fig. 3A, see Methods). Because measurements of individual antileukemic immune conditions are not available, we investigate three different approaches (fitting strategy I to III) for defining the relevant immune parameters *K*_*z*_ and *p*_*z*_ and compare the corresponding simulation results with the clinical data.

In *fitting strategy I*, we consider a generic immune system configuration with *identical* immune parameters *K*_*z*_ and *p*_*z*_ for all patients. Although the parameters can be adapted such that the overall rates and timings of recurrence meet the observed data (Fig. 3B), the model prediction fails on the individual level. The recurrence behavior could only be predicted correctly for 8 of 21 patients (Fig. 3D). This finding strongly argues in favor of patient-specific immune parameters.

In *fitting strategy II*, we estimate *patient-specific* immune parameter values by fitting the immune model to the pre-cessation BCR-ABL time courses. However, the immune system parameters *K*_*z*_ and *p*_*z*_ are not uniquely identifiable and vary over a broad range of possible values. Moreover, no statistically significant difference between recurring and non-recurring patients with respect to the fitted immune parameter values has been observed (Fig. S3). While model simulations show comparable recurrences on the population level, the individual prediction of recurrence behavior still fails (Fig. 3B/E).

In *fitting strategy III*, we demonstrate that *patient-specific* immune parameter values are sufficient to consistently explain the clinical data, although these parameters can only be estimated by fitting the immune model to the whole individual BCR-ABL time courses, including the post-cessation data. The model correctly describes the behavior on the population level (Fig. 3B), as well as on the individual patients (Fig. 3F-H). The patient-specific parameter estimates and the corresponding simulated BCR-ABL curves for all patients are presented in Fig. S4 and Table S1.

Having a closer look at the individually estimated parameters of the immune model using fitting strategy III, we observed that non-recurring patients show a significant higher immune proliferation rate *p*_*z*_ than recurring patients (p=0.02, Fig. 4A). We found no difference for the distribution of the immune inhibition constant *K*_*z*_ (p=0.57, Fig. 4B). To analyze the relationship between both immune parameters *K*_*z*_ and *p*_*z*_ in more detail, we did not only focus on the single best fit obtained from the parameter estimation routine (see Supplementary Materials) but had a concurrent view on different parameter combinations that all provide suitable fits for each patient. This reveals distinct parameter regions for recurring and non-recurring patients (Fig. 4F). A comparison of the estimated non-immunological parameters between recurring and non-recurring patients showed no statistically significant differences for the distributions of *p*_*XY*_ (Fig. 4C, p=0.09), *p*_*YX*_ (Fig. 4E, p=0.07) and *TKI* (Fig. 4D, p=0.89).

**Fig. 4.**
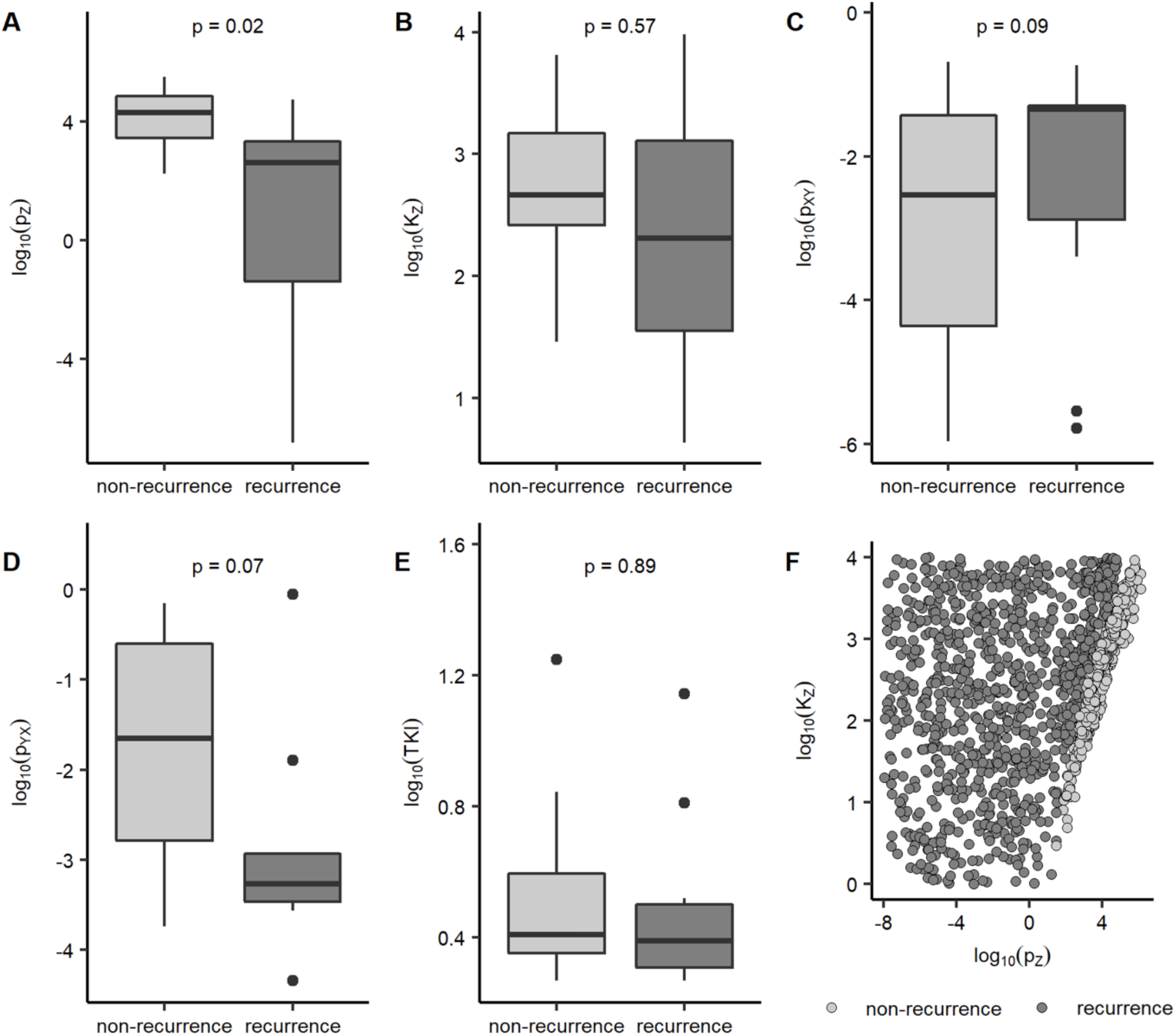
Comparison of estimated immune model parameters for recurring/non-recurring patients. **A-E** Boxplots of the best estimates for the immune model parameters *p*_*z*_, *K*_*z*_, *p*_*XY*_, *p*_*YX*_ and *TKI* obtained by using fitting strategy III, shown separately for the groups of recurring (dark grey) and non-recurring (light grey) patients. P-values of the corresponding Kolmogorov-Smirnov tests are indicated above. **F**: Scatter plot for the same immune parameters (*p*_*z*_, *K*_*z*_) for recurring (dark grey) and non-recurring patients (light grey). For this subfigure, 100 optimal parameter estimations (obtained from the ABC based optimization routine) of each patient using the immune model and fitting strategy III are shown.

From these results, we conclude that an individual immunological component (or another TKI-independent anti-leukemic effect) is necessary but also sufficient to quantitatively explain the individual BCR-ABL time courses of CML patients before *and* after stopping the TKI treatment. Our results suggest that for a particular patient no prospective prediction of the molecular risk of recurrence after TKI stop is possible based on the BCR-ABL dynamics under constant TKI treatment alone.

### Individual recurrence classification based on an “immunological landscape”

A detailed mathematical analysis of the calculated parameter values for the immune model estimated by fitting strategy III (see Supplementary text) suggests that the available patients can be grouped in three general classes that correspond to the underlying attractor landscape of the ODE model. It is the individual configuration of the leukemic and immunological parameters for each patient, which determines the outcomes (i.e. steady states) that can possibly be achieved after treatment cessation. In particular, each patient is characterized by the presence or absence of a *recurrence steady state E*_*H*_(Y ≈ K_y_), a *remission steady state E*_*L*_ with an immunological control of a residual leukemic disease (0<Y≪K_y_) or a *cure steady state E*_0_ in which the leukemic cells are completely eradicated (Y ≈ 0). The three classes are:

- Class A: For certain parameter configurations, the attractor landscape has only the stable recurrence steady state *E*_*H*_. This means that the patient will always present with recurring disease after treatment cessation due to an *insufficient immune response*, as long as the CML is not completely eradicated, irrespective of the degree of tumor load reduction. The corresponding attractor landscape is visualized in Figure 5A and depicts the recurrence behaviour depending on the number of immune cells and leukemic cells at treatment cessation. According to our estimates, 6 out of 21 patients fall into class A and ultimately present with recurring disease after treatment cessation (example in Figure 5B).
- Class B: For other parameter configurations, the attractor landscape has two stable steady states: the recurrence steady state *E*_*H*_ and the remission steady state *E*_*L*_. In this case there is a distinct remission level of BCR-ABL abundance, below which a *sufficient and strong* immune system can further diminish the leukaemia without TKI support. This also includes the possibility of a complete cure (Y ≈ 0). The corresponding attractor landscape indicates these two basins of attraction and is visualized in Figure 5C. We estimate 8 out of 21 patients in this class, which all maintain TFR (example in Figure 5D).
- Class C: The third class has the same stable steady states as class B, but in this case a small disturbance from the cure steady state *E*_0_ leads to the attraction basin of the recurrence steady state *E*_*H*_ instead of the remission steady state *E*_*L*_. Only for a small range of CML abundance and a sufficiently high level of immune cells, the immune system is appropriately activated to keep the leukaemia under sustained control. Figure 5E illustrates this control region as an isolated attractor basin. In the ideal case, the TKI therapy only reduces the leukemic load to a level that is sufficient to still activate the immune system to achieve this balance. However, if TKI treatment reduces tumor load to a very deep level, the CML regrows after therapy cessation as the immune response was also reduced too much. This represents a CML patient which may potentially achieve TFR but has a *sufficient but weak* immune response. We estimate that seven out of 21 patients fall in this class, of which four have a recurrence and three remain in TFR (two examples in Figures 5/G).

**Fig. 5.**
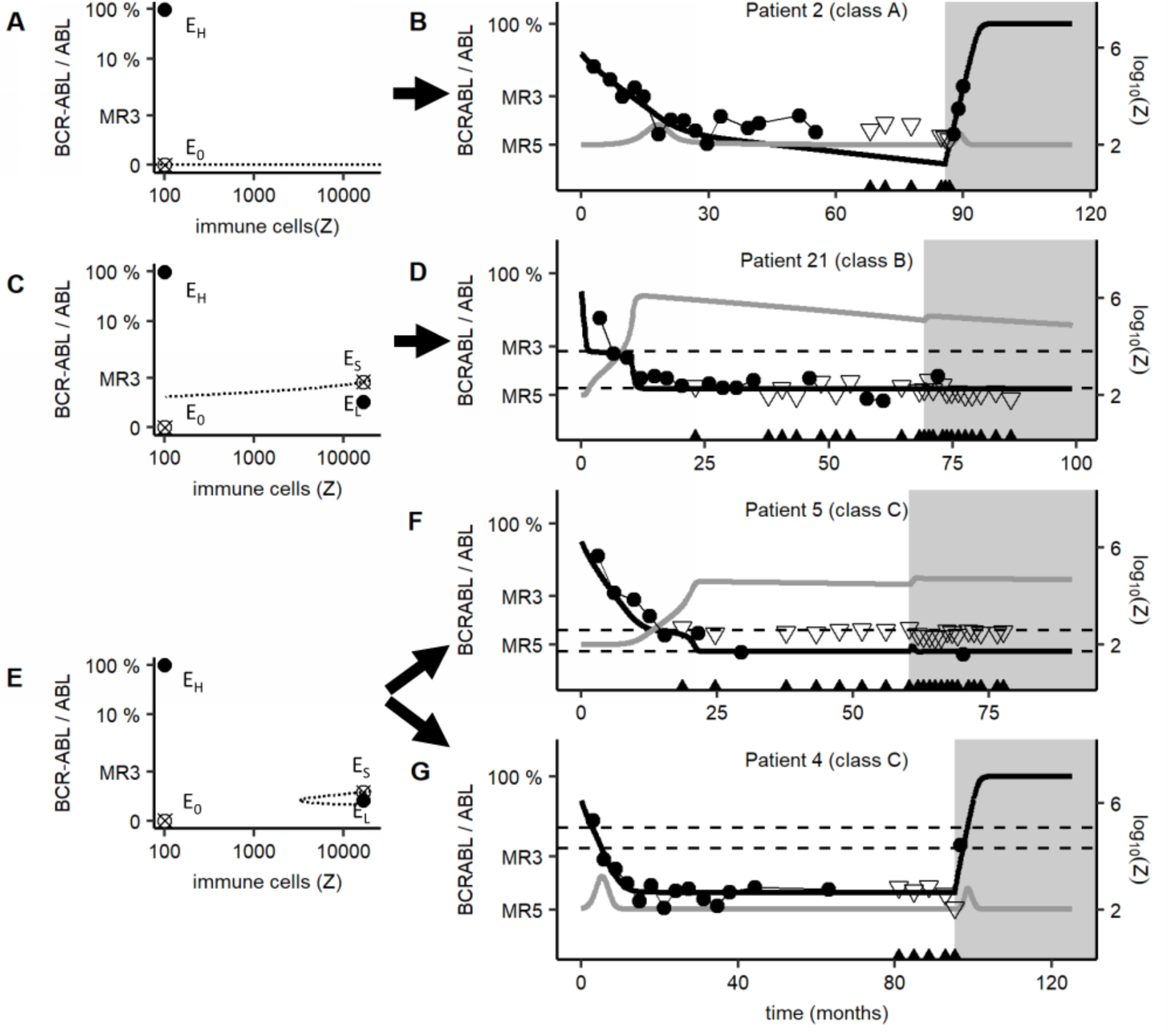
Model attractor landscapes and possible clinical scenarios. Representation of the attractor landscapes with corresponding clinical data and simulation results of the immune model using fitting strategy III for patients of the three classes. The phase space shown on the left side is restricted to the two-dimensional plane in which the BCR-ABL/ABL level and the number of immune cells are shown. Stable steady states are represented by solid circles (•) while saddle steady states are represented by crossed circles (⊗). The y-axis is set to a nonlinear scale via a root transformation. Clinical data and simulation corresponding to each phase portrait are shown on the right side: BCR-ABL measurements are shown as dots, triangles indicate the lower quantification limit for undetectable BCR-ABL levels. The grey area indicates the time after treatment cessation. If an immune window (i.e. a range of leukemic cells in which the expansion of the immune cell population is stimulated sufficiently) is existent, it is depicted by the area between both horizontal dashed lines. Calculated BCR-ABL values (thick black line) and corresponding relative number of immune cells (grey line) are shown on a logarithmic scale. (**A**) For patients in class A, there is only one basin of attraction with the recurrence steady state *E*_*H*_, leading to a recurrence after treatment cessation (**B**). (**C**) Class B patients present basins of attraction for the recurrence steady state *E*_*H*_ and remission steady state *E*_*L*_. The separatrix between the basins of attraction is represented by dashed lines passing through a saddle-point *E*_*S*_. (**D**) The corresponding, representative patient stays in remission after treatment cessation. (**E**) Patients of class C also present two basins of attraction but may show a recurrence after stopping TKI treatment or not (**F/G**).

For completeness, there is a fourth class, in which only a cure steady state exists. In this case, CML would not develop at all due to a strongly suppressive immune system. Naturally, those individuals do not appear in the patient cohort at all.

### Treatment optimization informed by the immunological configuration

We learnt that the immunological configuration of each patient determines the available steady states that can be reached using TKI treatment. It appears, that patients in class A can only stop TKI treatment in the case that the disease is completely eradicated. This would require a median treatment time of 29 years in our simulations and was not achieved in any of the considered patients. However, even if treatment cessation is no option for these patients, our previously published results suggests that TKI dose reduction could be considered as a treatment alternative (9).

From a perspective of treatment optimization, the patients in classes B and C are most interesting as they confer an *immune window*, in which a TKI-based reduction of the leukemic cells is able to sufficiently stimulate the expansion of the immune cell population (Fig. S5, Supplementary text). Patients in class B are characterized by an immunological response that is sufficient to control the leukemia once the leukemic load has initially been reduced below a certain threshold. This remission allows for an activation of the immune system to further control the leukemia eradication even in the absence of TKI treatment. It is essential that the initial remission and the immunological activation surpass a certain threshold, which is indicated by the line separating the different attractor basins in Figure 5C. Clinically, this can be achieved by a sufficiently long TKI therapy, although we predict that the respective 6 patients could all have already stopped their treatment much earlier as they actually did (compare Table S2).

In contrast to class B, patients in class C can also present with recurrence if a long TKI treatment is applied. Only in a narrow region of CML abundance the immune system is sufficiently stimulated: if leukemic load is too high, the immunological component is still suppressed, while for too low levels the stimulation is not strong enough. In this respect, TFR can only be achieved if treatment keeps the patient within his immune window for a sufficient time thereby supporting the adequate proliferation of immune cells, such that the patient reaches the attraction basin of the remission steady state *E*_*L*_ (Fig. 6A/B). If the treatment intensity is too high or the treatment duration is too long, this might lead to an “overtreatment” where the inherent immunological defense is not quickly and sufficiently activated to control a recurrence once TKI is stopped (Fig. 6C-F). It can be shown theoretically that an adjustment of the necessary balance between leukemia abundance and immunological activation can be achieved by detailed assessment of both critical cell populations and a titrated and narrowly adapted TKI administration (Fig. S6/7).

**Fig. 6.**
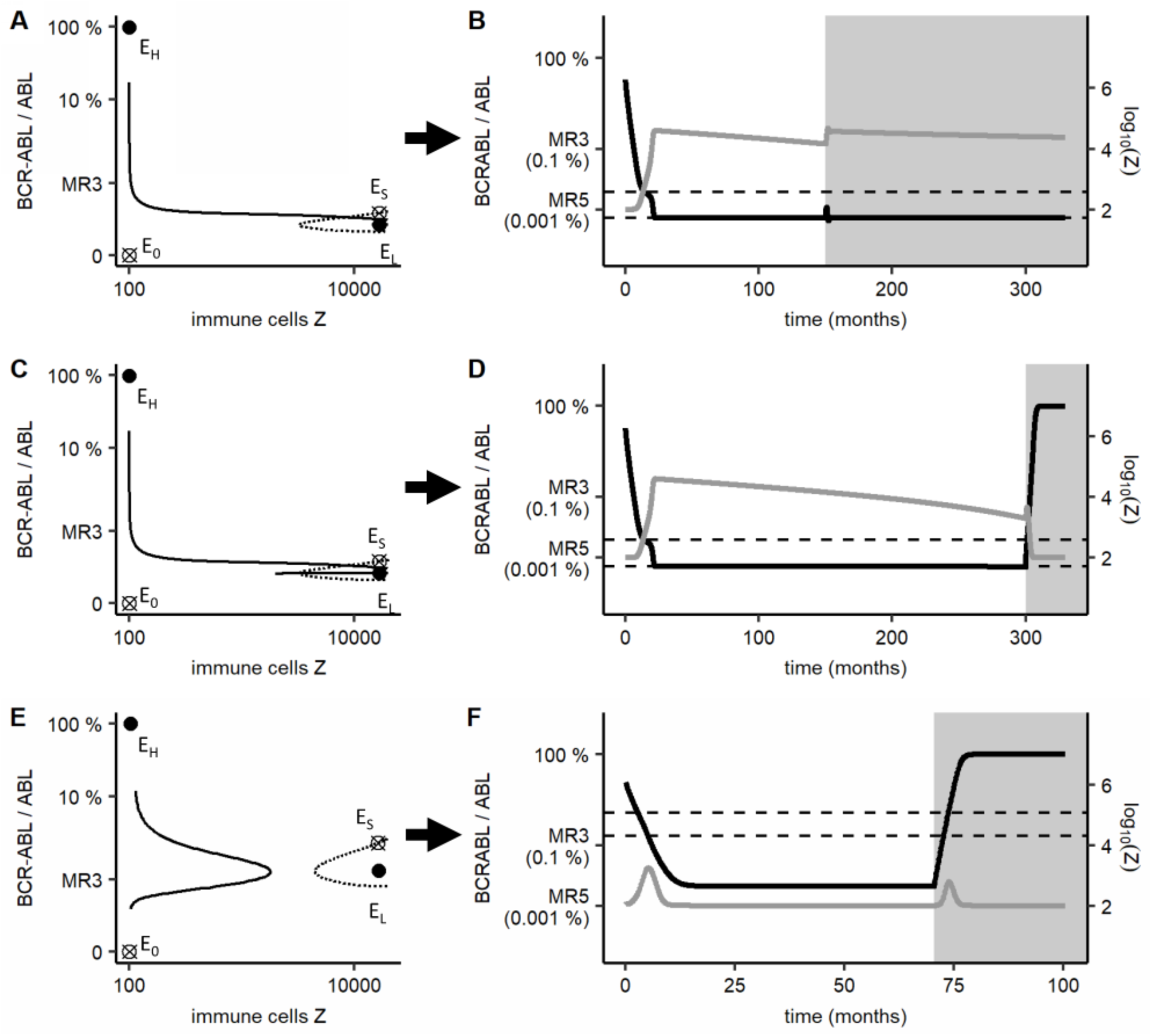
Predicted recurrence behavior of patients with a weak immune response (class C) Prediction of the recurrence behavior for patients with a weak immune response (i.e. class C) obtained by using the immune model and fitting strategy III. In comparison to Fig. 5, also the course of the number of leukemic cells and immune cells before stopping treatment (solid line) is shown within the phase space. If TKI treatment “drives” the system within the isolated basin of the remission steady state *E*_*L*_ (**A**), the simulated patient achieves a TFR (**B**). Simulating an extended treatment for this patient leads to another crossing of the separatrix curve (**C**) and thus, a recurrence after stopping treatment (**D**). A class C patient who does not cross the separatrix curve and thus, does not reach the remission basin, is shown in the last phase portrait (**E**). This leads to disease recurrence for this patient (**F**) independent of the applied treatment time.

### TKI dose alteration informs molecular recurrence after treatment cessation

Detailed information about a patient’s response to TKI treatment cessation (according to fitting strategy III) can only be obtained if the complete data (including post-cessation measurements) is available. Thus, this approach can obviously not serve as a prediction strategy *before* therapy stop. However, we show in the following that response to dose reduction – prior to therapy stop – will also depend on the immunological landscape and is, therefore, likely to provide important information about the disease dynamics after treatment cessation. Both, clinical and modeling evidence support the strategy to use information from intermediate dose reduction as this appears as a safe treatment option for almost all well responding CML patients (9,38).

Specifically, individual fits for all patients according to the immune model and fitting strategy III allow to mathematically simulate how the patients would have responded if they were treated with a reduced TKI dose instead of stopping TKI completely. We use these model simulations to derive information about the predicted BCR-ABL ratio during a 50% dose reduction within a 12-month period. Figure 7A/B illustrates two typical time courses.

**Fig. 7.**
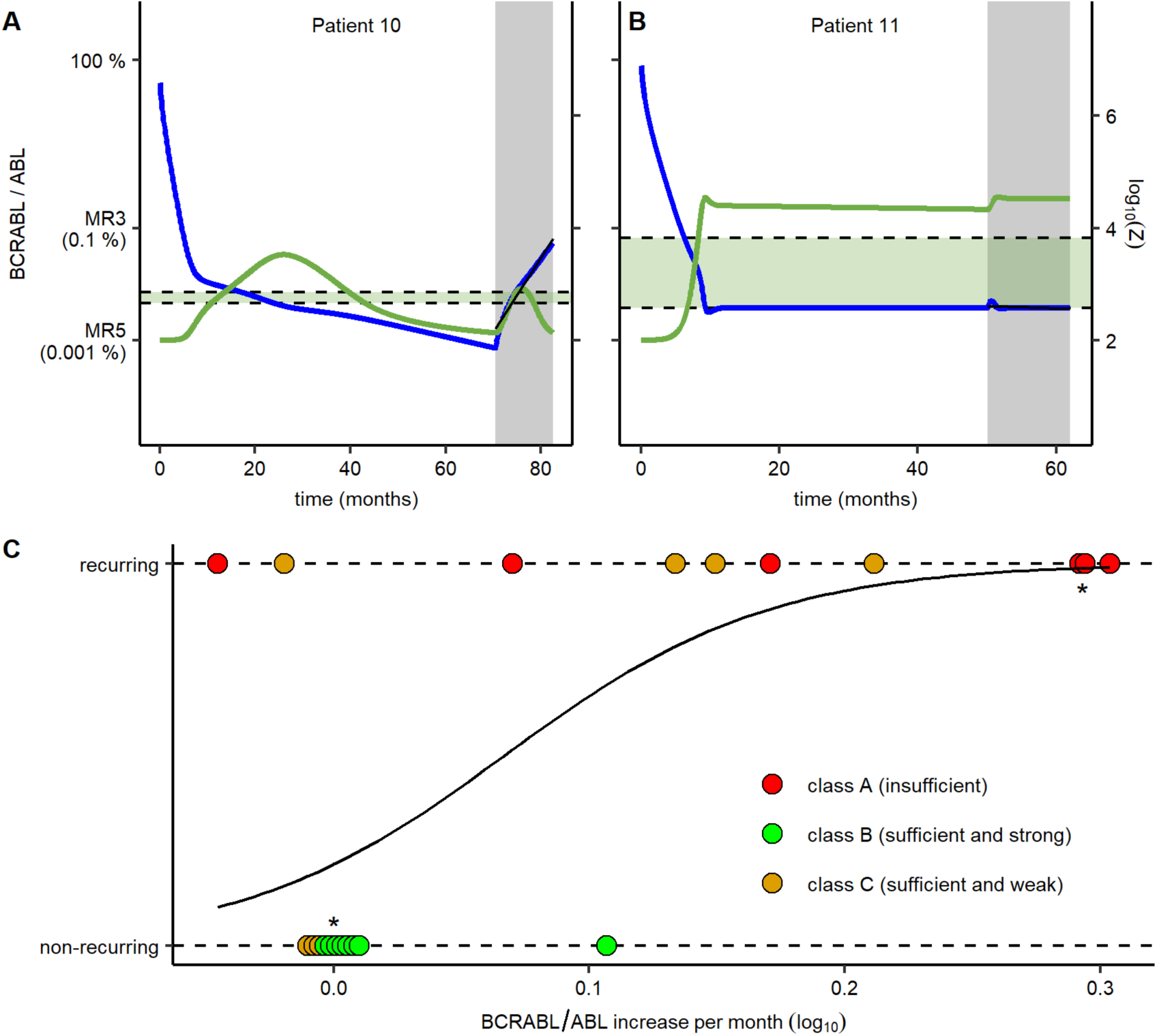
Individual response to dose reduction and association with the final remission state. **A/B:** Representative time courses of a simulated 12 months dose reduction period (grey background) for a recurring patient (**A**) and a non-recurring patient (**B**) with the corresponding treatment intensity using the immune model and fitting strategy III. The slope of the dose reduction period (thin black line) is calculated using a linear regression model. Computed BCR-ABL values (blue line) and corresponding relative number of immune cells (green line) are shown on a logarithmic scale. The light-grey area indicates the dose reduction period. The immune window is depicted as green background. **C**: Depiction of the logistic regression model for the final remission status of the 21 patients in the clinical trials versus their calculated slope within the dose reduction period. Also, the association of the patients to the three classes is shown (class A: red, class B: green, class C: brown). Overlapping points are stacked horizontally (*).

Quantitatively, we apply a linear regression model to estimate the individual BCR-ABL response slopes during this 12 months of dose reduction period. Depicting the final remission status after treatment cessation versus this slope (Figure 7C) indicates that recurring patients are predicted to be present with higher (positive) slopes of the BCR-ABL ratio during the dose reduction period. Moreover, complementing this plot with the association of each patient with its particular response class A, B, or C, we observe that class A patients have higher positive slopes and always have a recurrence, while most of class B patients show constant BCR-ABL levels, therefore, staying in TFR. Class C patients show both, constant or increasing BCR-ABL levels. However, higher positive slopes are more often observed in recurring patients. Therefore, we conclude that patients with pronounced increases in BCR-ABL levels after dose reduction correspond to either class A or C and should not stop TKI treatment (i.e. high risk for molecular recurrence), while patients with moderate slopes are predominantly found in class B and did most likely establish immune control, resulting in a higher chance to remain in TFR after therapy cessation.

## Discussion

Here we present a mathematical CML model that explicitly includes an immunological component and apply it to describe the therapy response and recurrence behavior of a cohort of 21 CML patients with detailed BCR-ABL follow-up over their whole patient history. We demonstrate that an anti-leukemic (most likely immunological) mechanism is necessary to account for a TKI-independent disease control, which prevents molecular recurrence emerging from residual leukemic cell levels after TKI cessation. Without such a mechanism, a long-term TFR can only be achieved if a complete eradication of leukemic cells is assumed. However, the presence of detectable MRD levels in many patients after therapy cessation (12) is not consistent with this assumption, which strongly suggests an additional control instance, which others (23,24,39,40) and we (35) interpret as a set of immunological factors. Including these aspects into our modeling approach, the available clinical data can be sufficiently described on the level of individual patients.

Based on our simulation results we classify patients into three different groups regarding their predicted immune system configuration (“immunological landscape”): i.e. *insufficient immune response* (class A), *sufficient and strong immune response* (class B) and *sufficient but weak immune response* (class C). Class A patients are not able to control residual leukemic cells and would always present with CML recurrence as long as the disease is not completely eradicated. Consistent with the results of Horn et al. (41), this is only accomplished on very long timescales and would, therefore, result in a lifelong therapy for most affected patients. However, as we suggested earlier, those patients might be eligible for substantial TKI dose reductions during maintenance therapy (9). In contrast, class B patients with a strong immune response are able to control the leukemia once the leukemic load has been reduced below a certain threshold and thus, require only a minimal treatment time (less than 5 years for the studied patients, see Table S2) to achieve TFR. For class C patients with a weak immune response, our model predicts that TFR achievement depends on an optimal balance between leukemia abundance and immunological activation before treatment cessation and could be accomplished by a narrowly adapted TKI administration.

We show that the information required to classify the patients according to their immune response and to predict their recurrence behavior cannot be obtained from BCR-ABL measurements *before* treatment cessation only. A different fitting strategy (III) assessing also BCR-ABL measurements after treatment cessation shows that the BCR-ABL changes from this system perturbation (i.e. TKI stop) yields the necessary information. Interestingly, our simulation results demonstrate that also a less drastic system perturbation, i.e. a TKI dose reduction, would provide similar information and can be used to predict the individual outcome after treatment cessation. The feasibility of such an approach has been recently substantiated by the DESTINY trial (NCT01804985), which evaluated a beneficial effect of a 12 month dose reduction treatment prior to TKI stop (38).

Direct measurements of the individual immune compartments and their activation states represent another road to better understand the configuration of the anti-leukemic immune response in CML patients. Several studies identified different immunological markers in CML patients that correlate with the probability of treatment-free remission after therapy cessation (23,24). Learning from the behavior of these populations under continuing TKI treatment and with lowered leukemic load could further contribute to identify a patient’s “immunological landscape” and be informative for the prediction of individual outcomes after treatment stopping. However, as it is not clear, which immunological subset provides the suggested observed anti-CML response, corresponding measurements are currently not feasible and strongly argue in favor of our indirect modeling approach suggesting to retrieve similar information from BCR-ABL dynamics after TKI dose reduction.

Our analysis is based on a rather small cohort of patients. Although our results do not depend on the study size, we can derive the strongest conclusions with respect to illustrating the conceptual approach of inferring immune responses from treatment alterations and demonstrating its predictive power. Our results are further based on a set of simplifications and assumptions. As such, we do not consider resistance mutations as almost no such events have been reported during TKI cessation studies in CML. Instead, it has been shown that different immune cell types are associated with recurrence behavior of CML patients (23,24). However, for simplicity, we restricted our analysis to an unified anti-leukemic immune compartment in the model and did not distinguish between different immune cell populations and interactions between them. Furthermore, the model is based on an interaction between leukemic and immune cells, in which the immune cell population is only activated for intermediate levels of leukemic burden, reflecting the assumption that immune cells are not efficiently activated for small numbers of leukemic cells and are additionally suppressed by high tumor load. Similar assumptions have been discussed recently (33) while we also illustrated the suitability of other mechanisms of interaction (35).

In summary, our results support the notion of immunological mechanisms as an important factor to determine the success of TFR in CML patients. Importantly, we show that besides the direct measurement of the immune response also system perturbations, like a TKI dose reduction, can (indirectly) provide information about the individual disease dynamics and, therefore, allow to predict the risk of CML recurrence for individual patients after TKI stop. Such personalized therapy decisions can increase life quality of the individual patients and additionally reduce treatment costs.

## Acknowledgement

We thank all patients and hospital staff for providing this valuable data for scientific assessment.

