## Supplementary Material for "Inferring immunological control mechanisms from TKI dose alterations in CML patients"

### Patient data

In total, 60 time courses of TKI-treated CML patients, for which TKI-therapy had been stopped as a clinical intervention, were provided by clinical partners. In particular, the patient data was obtained from

- the EURO-SKI trial (NCT01596114, 14 patients initially treated in Bordeaux, 2 patients initially treated in Poitiers, 5 patients initially treated in Helsinki, 5 patients initially treated in Mannheim)
- the STIM (NCT00478985, 3 patients initially treated in Bordeaux)
- STIM2 trials (NCT01343173, 11 patients initially treated in Bordeaux, 1 patient initially treated in Poitiers)
- local registries (1 patient initially treated in Bordeaux, 18 patients initially treated in Munich).

Informed written consent was obtained from each subject according to the local regulations of the participating centers.

Table S1 provides a detail account of the clinical trial/local registry of origin for all of the 21 patients that have been selected for the statistical data analysis.

### Calculation of the BCR-ABL/ABL ratio

To calculate the BCR-ABL Ratio on the decadic logarithm scale ($LRATIO$) in the peripheral blood, we use the following two assumptions: (I) We assume a maximum cell number (consisting of leukemic cells in Y and healthy cells) equal to the carrying capacity $K_y$. Thus, we calculate the number of healthy cells $y$ using the following equation: $y = (K_y - Y)$. (II) We assume that healthy cells express no BCR-ABL and twice as much ABL than leukemic cells.

Thus, we calculate the BCR-ABL/ABL ratio on a decadic logarithmic scale using:

$$LRATIO = log_{10}(\frac{Y}{Y + 2(K_y - Y)}) \quad (4)$$

### Derivation of the "immune window"

We define the immune window as the range of leukemic cells, for which the proliferation of immune cells $p_z * \frac{Y}{{K_z}^2+Y^2}$ exceeds the immune apoptosis rate $a$. With $Z \neq 0$, the following condition must be fulfilled:

$$p_z * \frac{Y}{{K_z}^2 + Y^2} > a \tag{5}$$

Solving this equation for $Y$, we obtain the following range for the immune window:

$$\frac{p_z}{2a} - \sqrt{\frac{{p_z}^2}{4a^2} - {K_z}^2} < Y < \frac{p_z}{2a} + \sqrt{\frac{{p_z}^2}{4a^2} - {K_z}^2} \tag{6}$$

Moreover, the immune window is only existent for patients with:

$$\frac{{P_z}^2}{4a^2} > {K_z}^2 \tag{7}$$

**Parameter estimation**

We use fixed values for the immune mediated killing rate $m$, the carrying capacity $K_Y$, immune cells natural influx $r_z$ and the immune cells apoptosis rate $a$. $r_z$ and $a$ are set to $200\ cells/month$ and $2.0\ month^{-1}$ , respectively, to ensure a low, ground state compartment size of 100 CML-specific immune cells. For the carrying capacity $K_Y$ of proliferating leukemic stem cells we assume a maximum number of 1 million cells (1–3) although our general results are invariant against a scaling of those quantities. We use a low value of $m = 1 * 10^{-4} cells^{-1}\ month^{-1}$ for the kill rate to ensure a minimal kill effect on leukemic cells for the ground state immune level.

Estimates for the treatment effect $TKI$ and for the proliferation rate $p_Y$ cannot be uniquely identified using pre-cessation data only as the initial BCR-ABL decline is determined by the net difference between those values (see equation (2)). We circumvent this limitation by estimating this proliferation rate from fitting the model to only post-cessation data of all recurring patients (i.e. $TKI$=0). In those cases, we assume no relevant immunological effect on the BCR-ABL kinetics during recurrence and set the corresponding parameter values $a, p_Z, r_Z, K_Z, m$ to zero. We obtained a mean proliferation rate $p_Y = 1.658\ month^{-1}$ which is used for all further simulations.

Using an Approximate Bayesian Computation approach (*R* version 3.4.4, adapted version of package *EasyABC* version 1.5), we calculate a set of estimations for the remaining free parameters (i.e. the transition rates $p_{XY}$ and $p_{YX}$, the TKI kill rate $TKI$, and the immune parameters $K_z$ and $p_z$) for each patient, of which we use the best fitting parameter combination from each set for further simulations. The initial size of proliferating leukemic cells $Y$ is obtained from the initial BCR-ABL ratio and the carrying capacity $K_Y$. If no BCR-ABL ratio on treatment start is available, we estimate the initial BCR-ABL ratio from the bi-exponential fit. Given a constant cell number in Y, the values for X and Z are initialized in a quasi-steady state condition. Kolmogorov-Smirnov tests are used to compare the distribution of the estimated parameters between recurring and non-recurring patient groups (R version 3.4.4).

**Calculation of steady states**

We observe three different steady states in our immune model after stopping treatment (i.e., equations (1-3) with $TKI = 0$): (i) recurrence and two remission steady states, namely (ii) complete eradication and (iii) immunological controlled remission.

***Eradication steady state***

The complete eradication or complete remission steady state is the trivial equilibrium $E_0 = (X_0, Y_0, Z_0)$ of system (1-3). It is defined by $Y_0 = 0$, $X_0 = 0$ and

$$Z_0 = \frac{r_Z}{a}. \tag{8}$$

***Immunological controlled remission and recurrence steady states***

Both the immunological controlled remission steady state $E_L = (X_L, Y_L, Z_L)$ and the recurrence steady state $E_L = (X_H, Y_H, Z_H)$ are characterized by non-zero levels of leukemic cells in X and Y. However, the former is defined by low levels while the latter is defined by high levels of leukemic cells, i.e., $X_L < X_H$ and $Y_L < Y_H$. The steady states are obtained as follows. Setting $dX/dt = 0$ and isolating $X$ we obtain

$$X = (p_{YX}/p_{XY})Y \tag{9}$$

Analogously, setting $dZ/dt = 0$ and isolating $Z$ we obtain

$$Z = \frac{r_Z(K_Z^2 + Y^2)}{a(K_Z^2 + Y^2) - p_Z Y} \tag{10}$$

Now, setting $dY/dt = 0$ and substituting the expressions of $X$ and $Y$ we obtain, after simplification, the following third-degree polynomial equation in $Y$:

$$-a\, p_Y Y^3 + \big(p_Y p_Z + K_Y(ap_Y - mr_Z)\big)Y^2 + -p_Y(aK_Z^2 + K_Y p_Z)Y + K_Y K_Z^2(ap_Y - mr_Z) = 0 \tag{11}$$

Using the same techniques as in our previous publication (4) one can show the following. If $ap_Y < mr_Z$ then such equation does not have a positive solution $Y$. In this case, both the remission and recurrence steady states do not exist.

On the contrary, if $ap_Y > mr_Z$ then there are two options, depending on the discriminant of such equation: either the equation has one or three positive solutions $Y$. We omit the formula of the discriminant for sake of brevity. If the equation has three positive solutions, we denote them as $Y_H > Y_S > Y_L$. Substituting these values in the expressions of $X$ and $Z$ above, we obtain three equilibrium points, the recurrence steady state $E_H$, the remission steady state $E_L$ and a third steady state denoted as $E_S = (X_S, Y_S, Z_S)$. For all parameter values used here, we obtained that $E_H$ and $E_L$ are stable, while $E_S$ is a saddle point. This case corresponds to the landscape attractors C and E in Fig. 5, where two basins of attraction are present. The distinction of such cases is made by assessing whether a system solution starting as small perturbation of $E_0$ will converge either to $E_L$ or to $E_H$. Finally, if the above polynomial equation has one positive solution $Y_H > 0$, then it defines the recurrence steady state $E_H$, while the remission steady state does not exist. In this case, we numerically verify that $E_H$ is stable. Thus, this case corresponds to the attractor landscape in Fig. 5A.

**Treatment optimization approach**

Taking into account the concept of an optimal window for immune activity, we model also an alternative, adaptive TKI treatment. In addition to using a fixed treatment intensity $TKI_0$, we also apply an alternative treatment strategy, which reduces the treatment kill rate $TKI$ when the leukemic cells level is reduced below the immune window. This strategy aims to keep the leukemic cells level within the immune window for a longer time in such way it prolongs the stimulation of the immune cell proliferation. Formally, this is incorporated in the model by multiplying $TKI_0$ with a sigmoid function, thereby obtaining:

$$TKI = TKI_0 * \frac{1}{1 + e^{-\frac{\frac{Y}{Y_{MIN}} - 1}{0.01}}} \tag{12}$$

### Supplementary figures

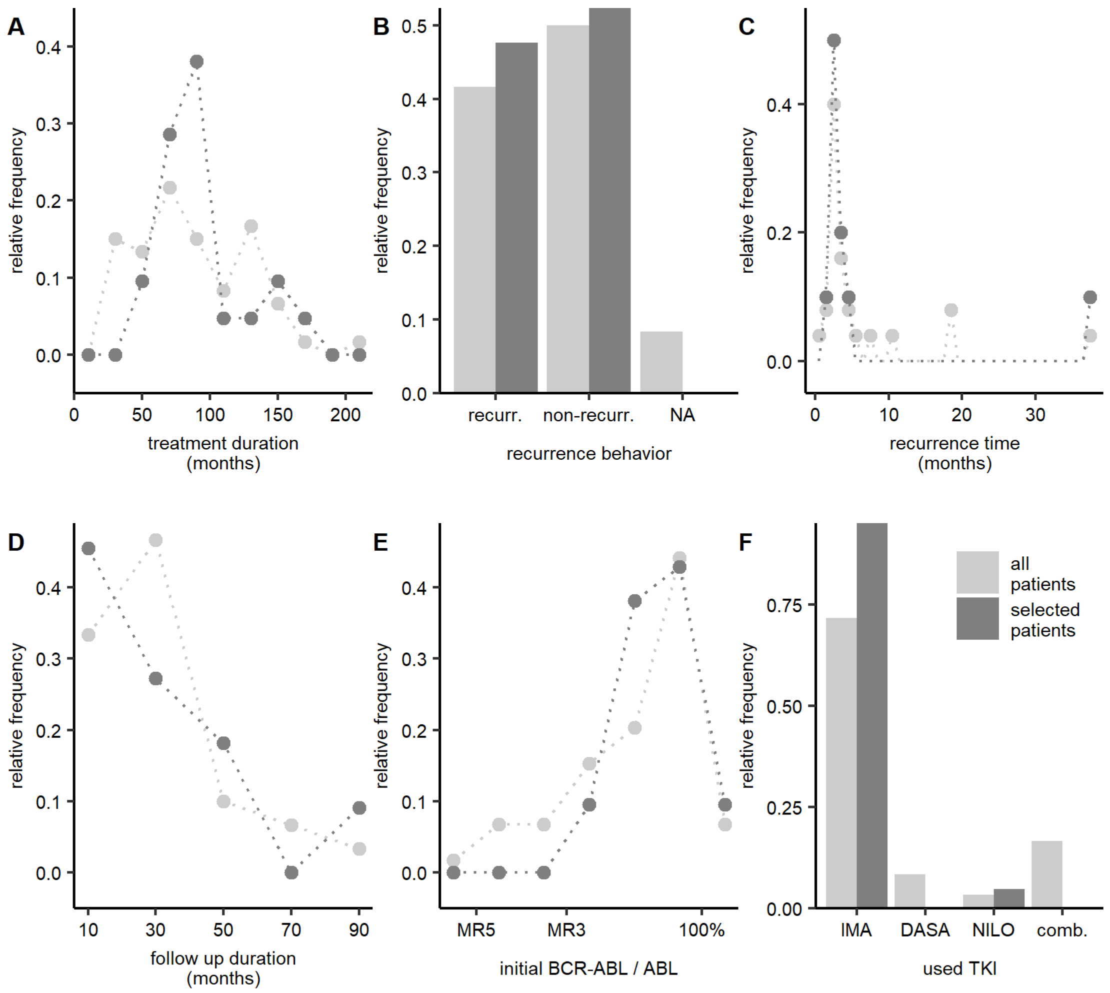


**Fig. S1 Clinical characteristics of original and selected patients**

**A-F**: Comparison of selected patients (dark grey, n =21) and all patients (light gray, n=60). The y axis depicts the relative frequency. **A**: Treatment duration until cessation. **B**: Relative frequency of patients recurring and non-recurring after stopping TKI treatment. NA indicates missing information about recurrence behavior of the patient. **C**: Time until occurrence of recurrence after stopping treatment. Recurrence is defined as loss of MMR (0.01% BCR-ABL) or begin of re-treatment. **D**: Follow up of BCR-ABL measurements of non-recurring patients after treatment cessation. **E**: Initial BCR-ABL levels. Patients with initial BCR-ABL/ABL measurements below MR3 has been excluded. **F**: Used TKI for treatment (DASA=Dasatinib, IMA=Imatinib, NILO=Nilotinib, comb.=combination therapy).

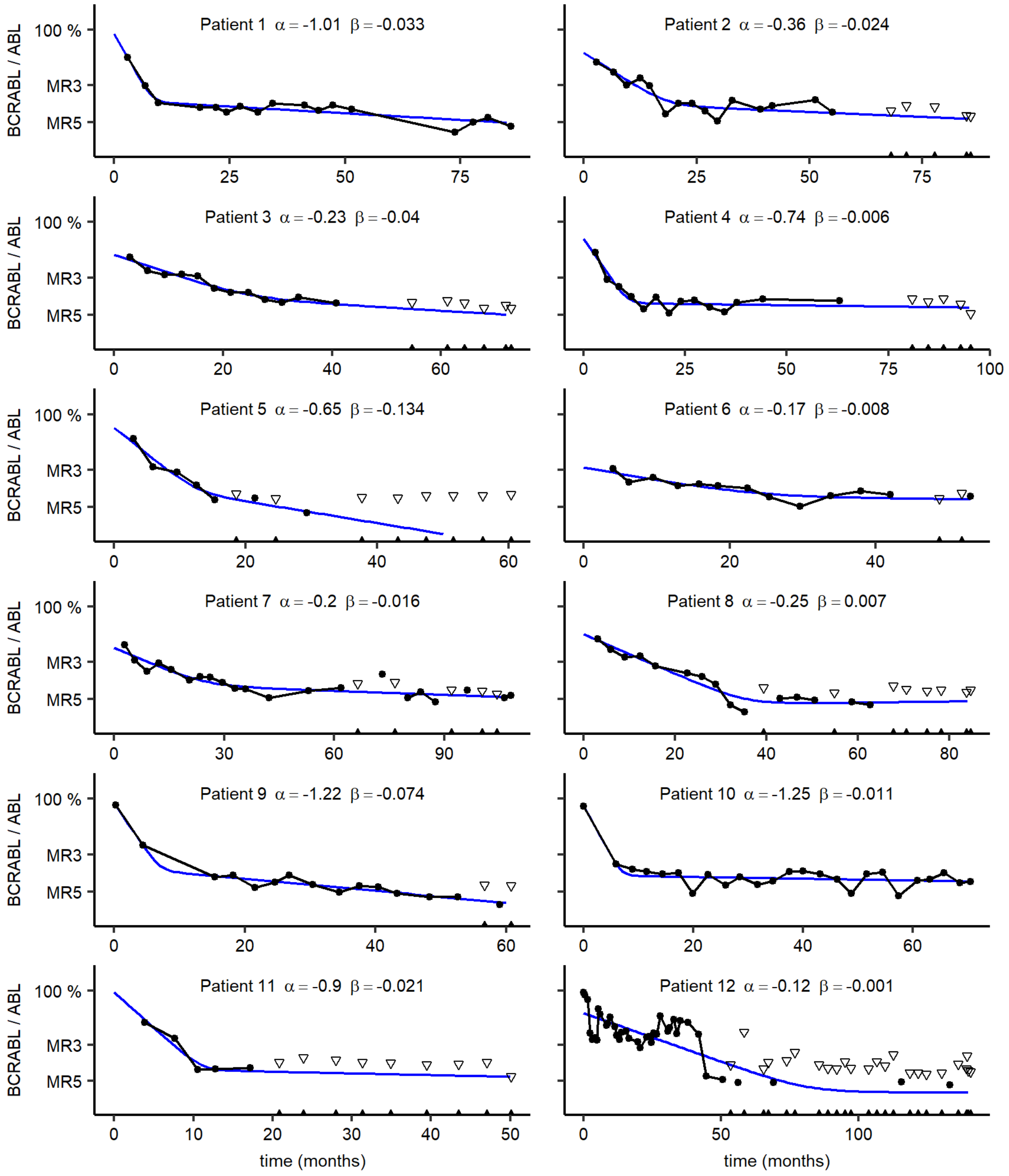


**Fig. S2 clinical data with corresponding bi-exponential fits (1)**

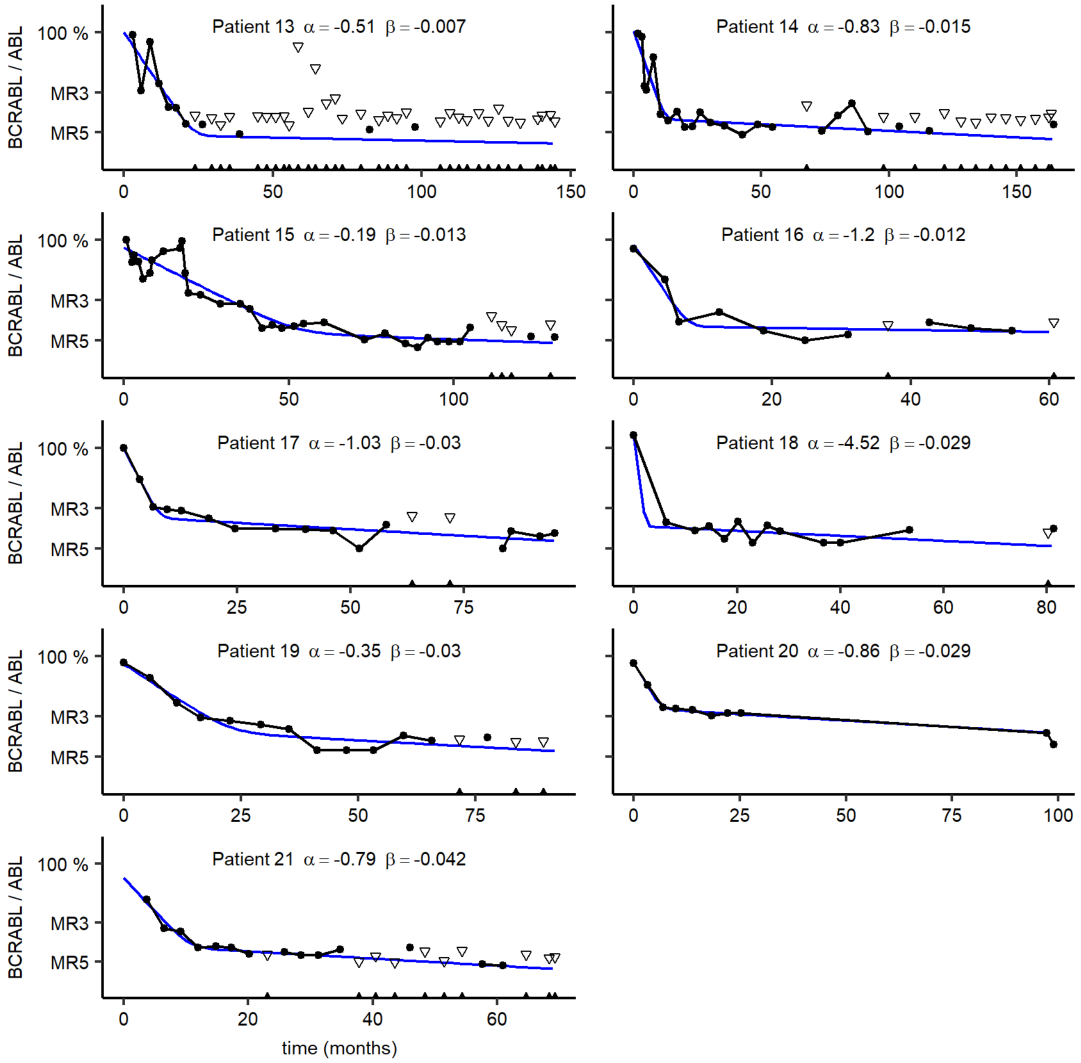


**Fig. S2 clinical data with corresponding bi-exponential fits (2)**
Clinical data and corresponding bi-exponential fits with corresponding α and β values for the 21 selected patients. BCR-ABL measurements are shown as black dots. For measurements with undetectable BCR-ABL levels, an upper bound is calculated based on the abundance of the reference gene. Therefore, the true BCR-ABL/ABL value is expected between this upper bound (open black triangle) and zero (filled triangles). The corresponding bi-exponential fit (blue line) consists of the initial fast decline α and the second, slow decline β. Only data until cessation is shown.

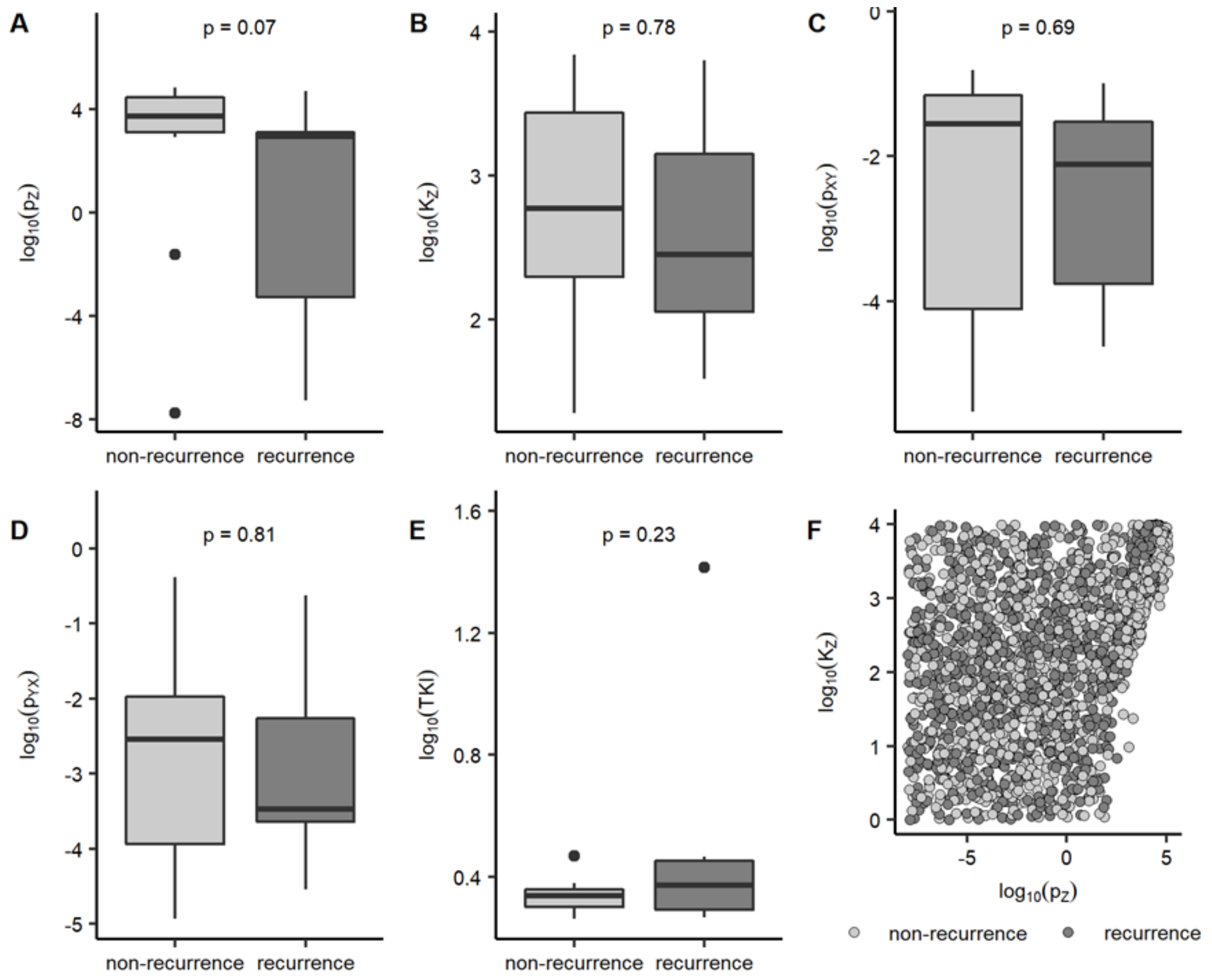


**Fig. S3 estimated model parameters by fitting data to measurements *before* cessation (fitting strategy II)**

**A-E**: Comparison of the parameter estimations for recurring (dark grey) and non-recurring (light grey) patients, estimated by fitting our immune system model to the initial BCR-ABL time courses *before* treatment cessation (fitting strategy II). P-values of the corresponding Kolmogorow-Smirnov tests are shown. **F**: Comparison of the best 100 estimations of immune system parameters $p_z$ and $K_z$ for recurring (dark grey) and non-recurring (light grey) patients.

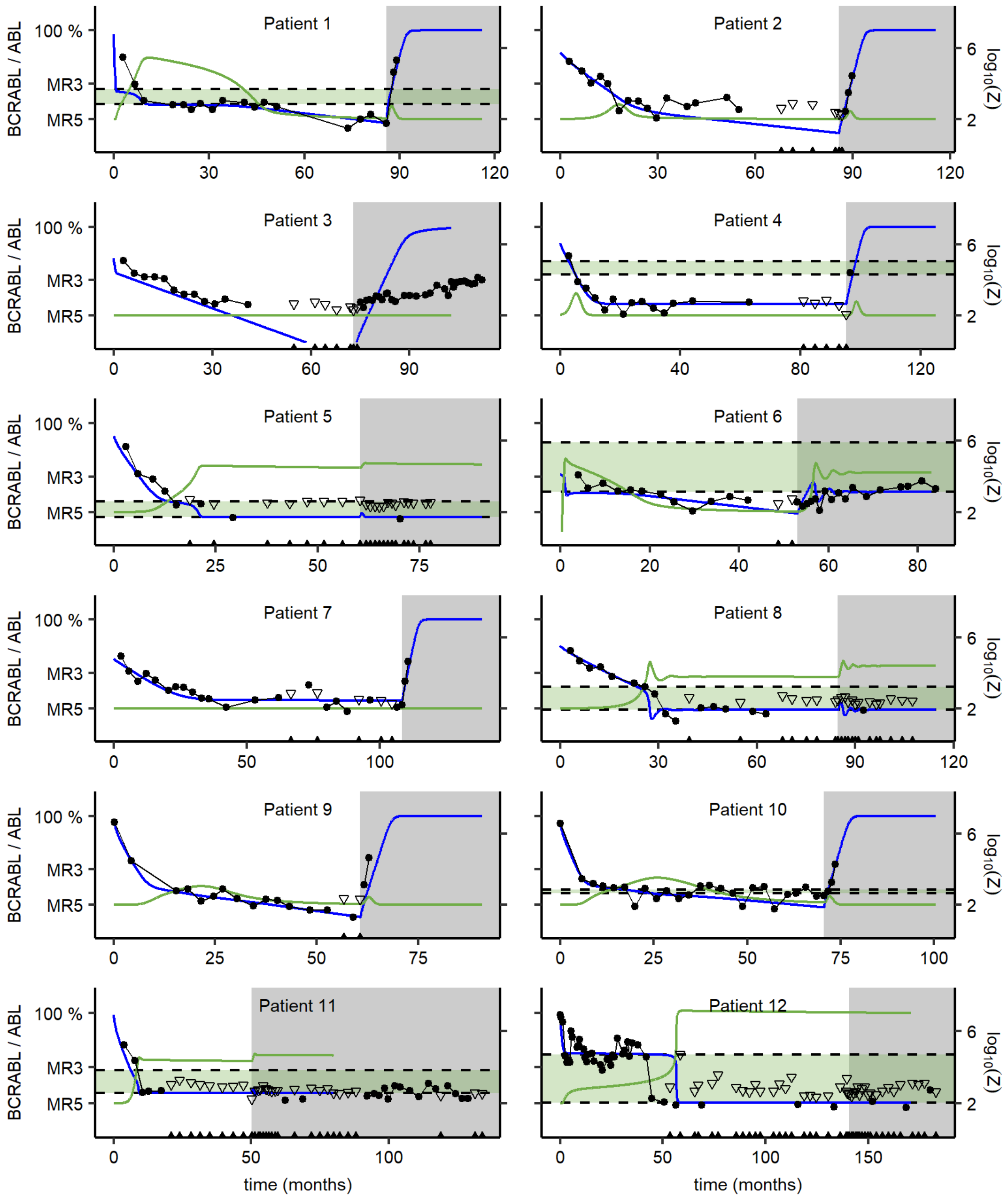


**Fig. S4: clinical data with corresponding immune system model simulations (1)**

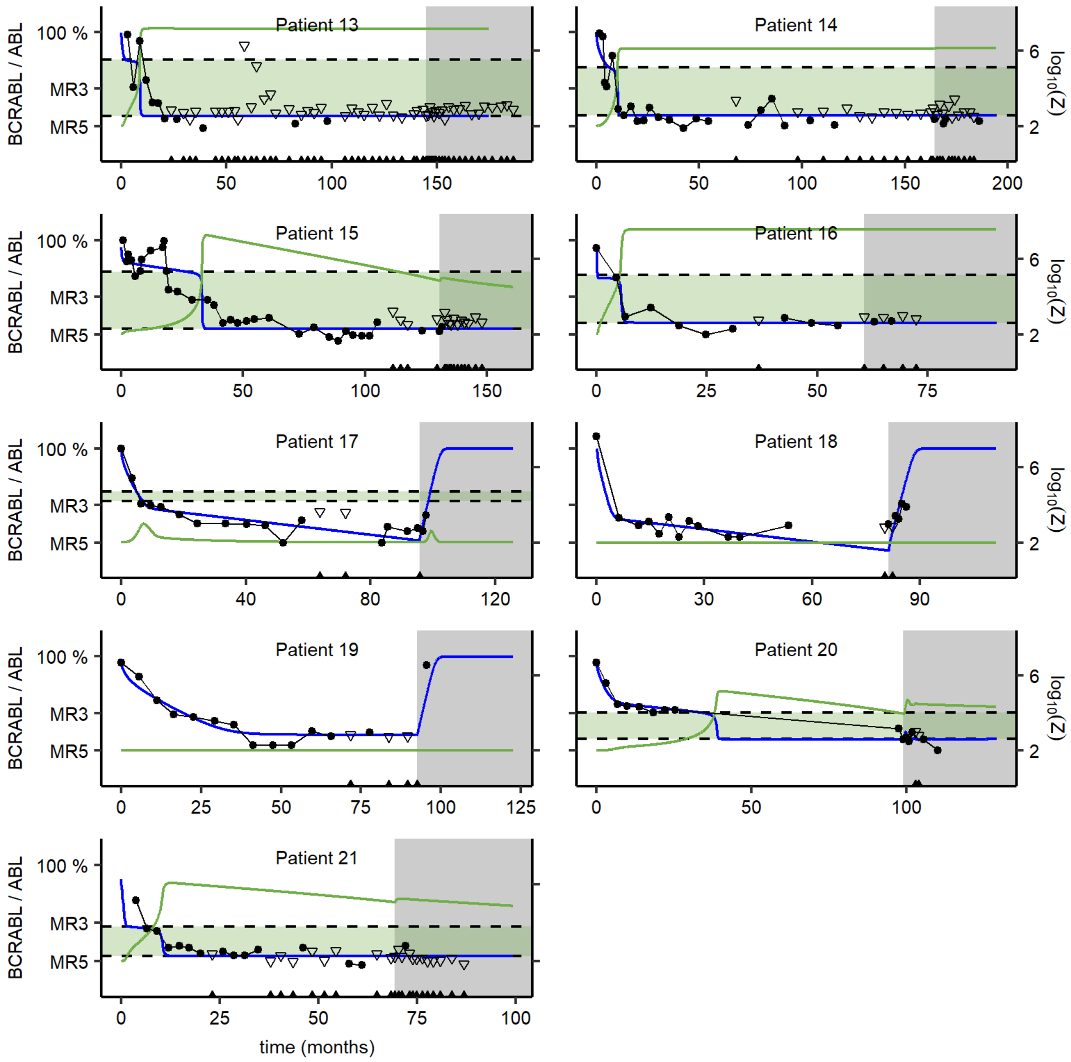


**Fig. S4: clinical data with corresponding immune system model simulations (2)**

Clinical data and corresponding immune model simulations of the 21 selected patients obtained by using fitting strategy III. BCR-ABL measurements are shown as black dots. For measurements with undetectable BCR-ABL levels, an upper bound is calculated based on the abundance of the reference gene. Therefore, the true BCR-ABL/ABL value is expected between this upper bound (open black triangle) and zero (filled triangles). The grey area indicates the time after treatment cessation. The immune window is depicted as green background. Calculated BCR-ABL values (blue line) and corresponding relative number of immune cells (green line) of the model on a logarithmic scale are shown. The following fixed parameter values are used for all patients: $K_Y$=1e+06, $m$=1e-04, $r_z$=200, $p_y$=1.658, $a$=2. The remaining, individual estimated parameter and the correspondence of the patients to the clinical trials and local registries can be obtained from Table S1.

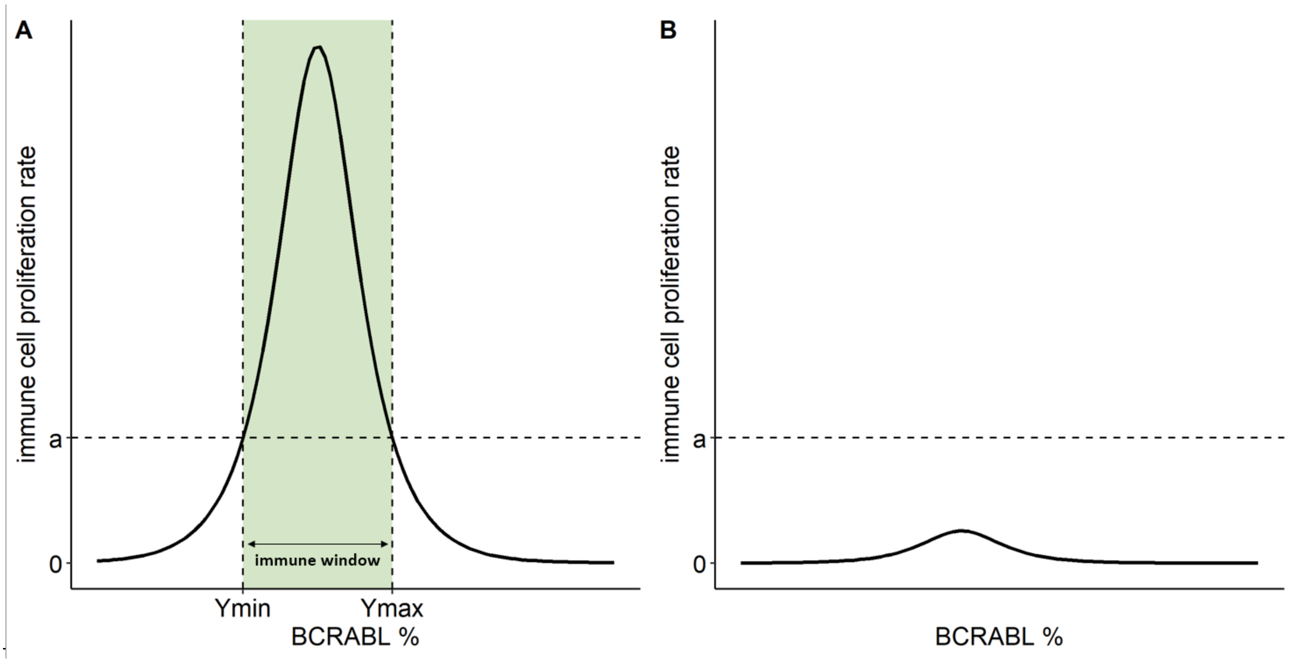


**Fig. S5 immune window**

We define the immune window (green area between the dashed lines) as the range of leukemic cells for which the immune cell proliferation rate exceeds the immune apoptosis rate $a$. The existence of the immune window depends on the individual values of the immunological parameter. Thus, it is existent in patients with an sufficient immune response (A) whereas patients with an insufficient immune response don't have an immune window (B).

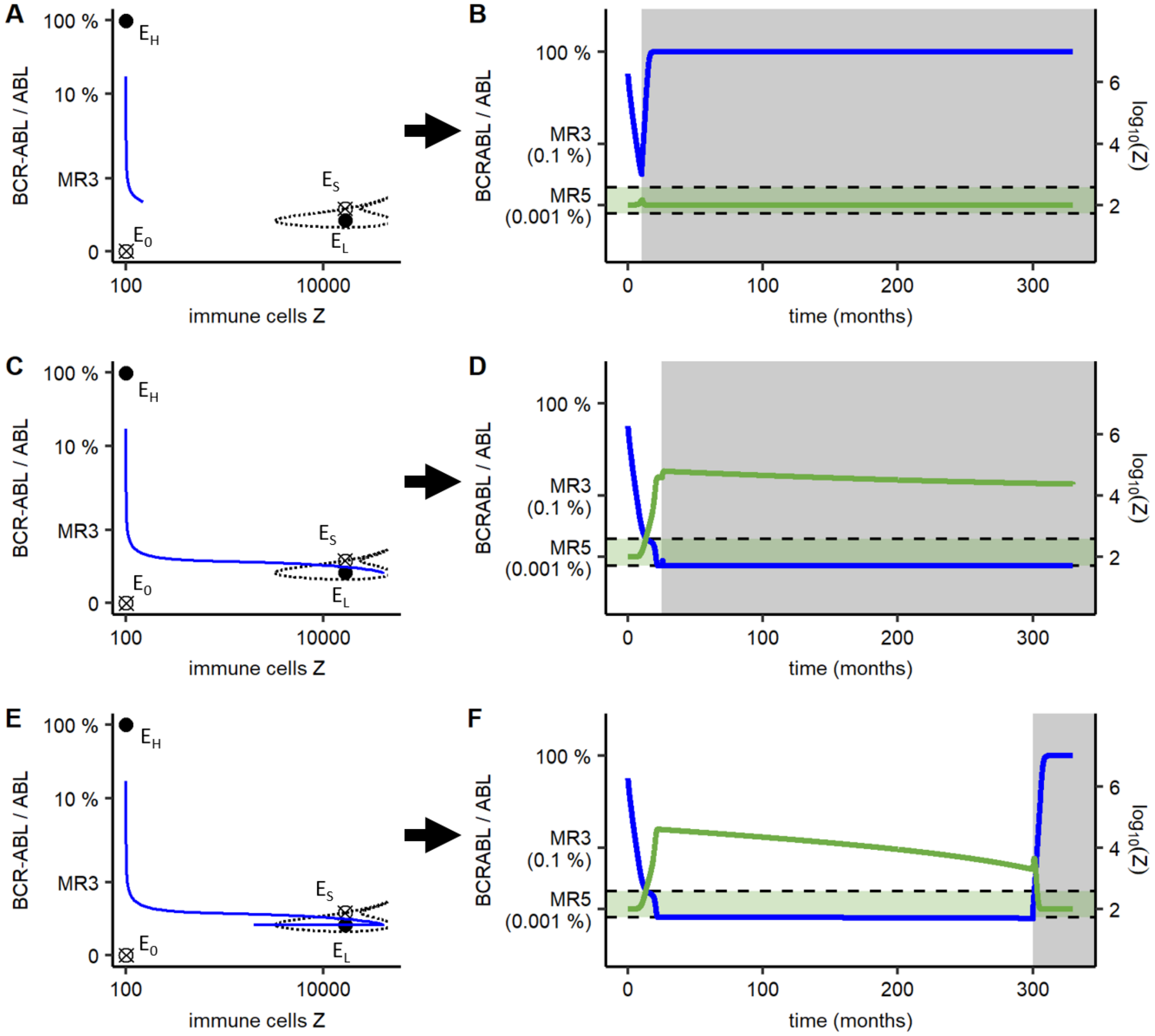


**Fig. S6 Recurrences of some patients with an estimated weak immune response (class C) can be prevented in our model predictions by applying an optimized treatment duration.** Representation of the attractor landscapes with corresponding clinical data and simulation results of patient 5 for different treatment times obtained by fitting the immune model using fitting strategy III. The phase space is shown on the left side (see description of Fig. 5 for details) together with the course of the number of leukemic cells and immune cells before stopping treatment (blue line). The time course of the BCR-ABL/ABL and immune cells including the time after stopping treatment corresponding to each phase portrait is shown on the right side. The grey area indicates the time after treatment cessation. The immune window is depicted as green background. Calculated BCR-ABL values (blue line) and corresponding relative number of immune cells (green line) on a logarithmic scale are shown.

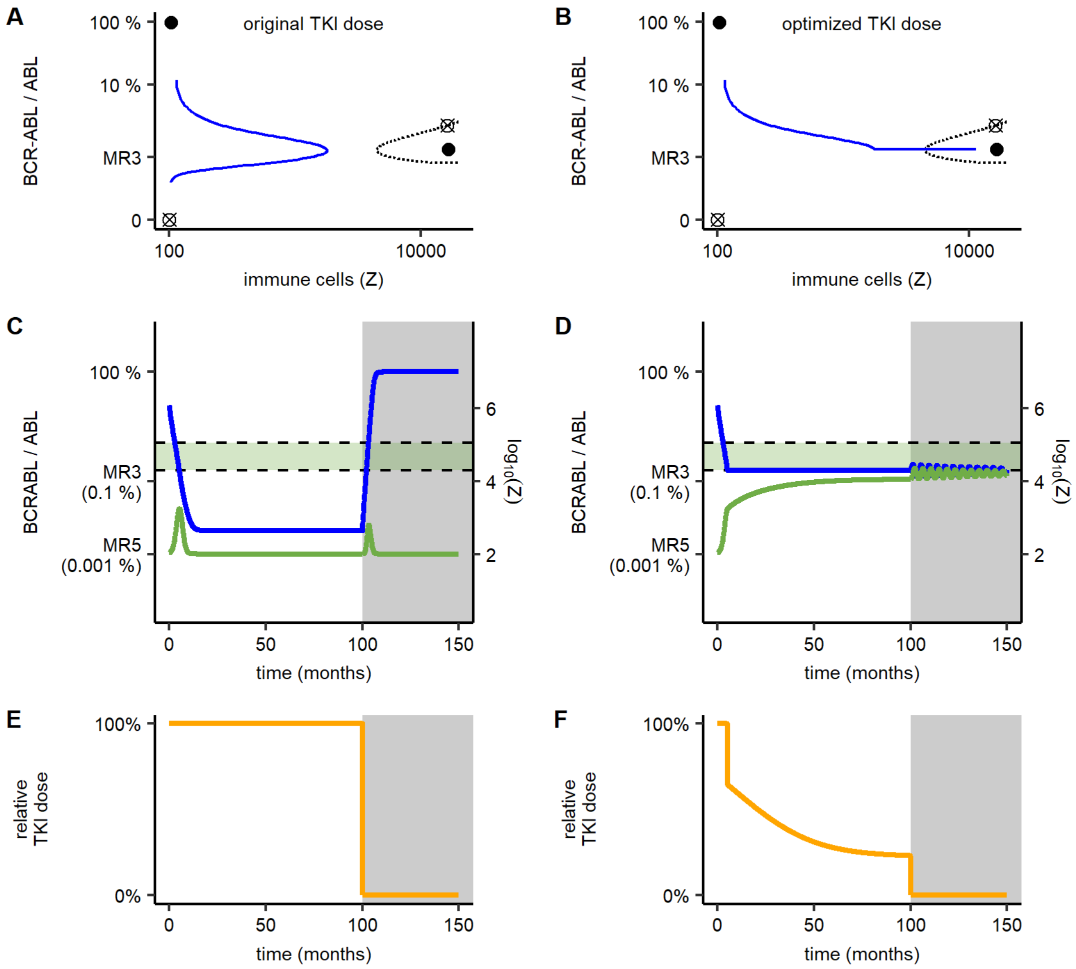


**Fig. S7 Recurrences of patients with an estimated weak immune response (class C) can be prevented in our model predictions by applying an optimized treatment intensity strategy.** The original treatment strategy using a constant value for the TKI intensity is shown on the left side. The optimized treatment approach using a treatment strategy which reduces the TKI dose when the lower limit of the immune window limit is reached is shown on the right side (see also Supplementary text). **A/B:** The phase portrait with the corresponding course of the leukemic cells and immune cells (see Fig. 5 for a detailed description). **C/D**: Depiction of the BCR-ABL/ABL and leukemic cell course. The grey area indicates the time after treatment cessation. The immune window is depicted as green background. Calculated BCR-ABL values (blue line) and corresponding relative number of immune cells (green line) of the model on a logarithmic scale are shown. **E**: The original treatment strategy applied in A/C. **F**: The optimized treatment approach using a treatment strategy which reduces the TKI dose when the lower limit of the immune window limit is reached (applied in B/D).

| Patient ID | $p_{YX}$ (months$^{-1}$) | $p_{XY}$ (months$^{-1}$) | TKI (months$^{-1}$) | $p_z$ (cells/months) | $K_z$ (cells) | cessation time (months) | clinical trial or local registry |
|---|---|---|---|---|---|---|---|
| **Patient 1** | 0.01 | 0.05 | 13.90 | 2210.00 | 362.00 | 85.90 | Bordeaux |
| **Patient 2** | 0.00 | 0.05 | 1.95 | 1940.00 | 508.00 | 85.70 | STIM2 (Bordeaux) |
| **Patient 3** | 0.88 | 0.18 | 6.48 | 0.00 | 4.29 | 72.90 | STIM2 (Bordeaux) |
| **Patient 4** | 0.00 | 0.00 | 2.31 | 53800.00 | 9570.00 | 95.20 | STIM2 (Bordeaux) |
| **Patient 5** | 0.00 | 0.01 | 2.38 | 176.00 | 28.70 | 60.30 | STIM2 (Bordeaux) |
| **Patient 6** | 0.70 | 0.21 | 2.07 | 311000.00 | 6480.00 | 53.00 | EUROSKI (Bordeaux) |
| **Patient 7** | 0.00 | 0.00 | 1.86 | 0.01 | 26.30 | 108.00 | EUROSKI (Bordeaux) |
| **Patient 8** | 0.00 | 0.00 | 1.86 | 696.00 | 74.70 | 84.70 | EUROSKI (Bordeaux) |
| **Patient 9** | 0.00 | 0.07 | 2.56 | 325.00 | 85.70 | 60.70 | EUROSKI (Bordeaux) |
| **Patient 10** | 0.00 | 0.04 | 2.77 | 473.00 | 115.00 | 70.50 | EUROSKI (Bordeaux) |
| **Patient 11** | 0.00 | 0.00 | 2.57 | 2820.00 | 318.00 | 50.10 | STIM (Bordeaux) |
| **Patient 12** | 0.04 | 0.00 | 4.30 | 19800.00 | 462.00 | 141.00 | Mannheim |
| **Patient 13** | 0.11 | 0.00 | 3.23 | 132000.00 | 2120.00 | 145.00 | Mannheim |
| **Patient 14** | 0.01 | 0.00 | 2.13 | 54000.00 | 1430.00 | 165.00 | Mannheim |
| **Patient 15** | 0.59 | 0.06 | 3.62 | 82600.00 | 1300.00 | 131.00 | Mannheim |
| **Patient 16** | 0.59 | 0.00 | 17.70 | 57600.00 | 1550.00 | 60.70 | Munich |
| **Patient 17** | 0.00 | 0.05 | 2.36 | 26800.00 | 5700.00 | 95.60 | Munich |
| **Patient 18** | 0.00 | 0.05 | 3.32 | 0.00 | 6.84 | 81.30 | Munich |
| **Patient 19** | 0.00 | 0.00 | 1.86 | 14.70 | 1740.00 | 92.60 | Munich |
| **Patient 20** | 0.01 | 0.04 | 2.38 | 4420.00 | 419.00 | 99.00 | Munich |
| **Patient 21** | 0.02 | 0.04 | 7.01 | 2530.00 | 212.00 | 69.30 | EUROSKI (Poitiers) |

**Table S1: individual parameter estimations and clinical trial/local registry of each patient**

Individual parameter estimations for the 21 patients obtained by using the immune model and fitting strategy III. The following fixed parameter values are used for all patients: $K_Y$=1e+06, $m$=1e-04, $r_z$=200, $p_y$=1.658, $a$=2. The corresponding clinical trial or local registry for the clinical data of each patient is shown.

| Patient ID | treatment duration (clinical data, months) | predicted minimal treatment time to achieve TFR (months) |
|---|---|---|
| **Patient 6** | 53.0 | 0.8 (1.5 %) |
| **Patient 12** | 140.8 | 56.5 (40.1 %) |
| **Patient 13** | 144.7 | 9.1 (6.3 %) |
| **Patient 14** | 164.6 | 10.4 (6.3 %) |
| **Patient 15** | 130.6 | 33.2 (25.4 %) |
| **Patient 16** | 60.7 | 5.7 (9.4 %) |
| **Patient 20** | 99.0 | 38.2 (38.6 %) |
| **Patient 21** | 69.3 | 9.9 (14.3 %) |

**Table S2 estimated minimal required treatment time of patients with a strong immune response (class B)**

Comparison of the treatment duration applied in the clinical trials with the predicted minimal treatment time required to achieve TFR in our simulations using the immune model and fitting strategy III. Only class B patients (i.e. with a strong immune response) are shown.
